# Genome-Encoded Cytoplasmic Double-Stranded RNAs, Found in C9ORF72 ALS-FTD Brain, Provoke Propagated Neuronal Death

**DOI:** 10.1101/248328

**Authors:** Steven Rodriguez, Benjamin R. Schrank, Asli Sahin, Hawra Al-Lawati, Isabel Costantino, Eric Benz, Darian Fard, Alefiya D. Albers, Luxiang Cao, Alexis C. Gomez, Elena Ratti, Merit Cudkowicz, Matthew P. Frosch, Michael Talkowski, Peter K. Sorger, Bradley T. Hyman, Mark W. Albers

## Abstract

Innate immune signaling activation and DNA damage are pathological hallmarks of aging that may herald multiple adult-onset neurodegenerative diseases. Here, we report that both cell autonomous and non-autonomous neuronal death are triggered by the production of cytoplasmic double-stranded RNA (cdsRNA) from a regulated, disarticulated transgene in the setting of type I interferon (IFN-I) signaling. CdsRNA is a pathogen associated molecular pattern that induces IFN-I in many cell types. Transfection of a dsRNA mimetic into cultured human neurons also induces IFN-I signaling and cell death in a dose-dependent manner. Direct relevance to human disease is found in neurons of ALS-FTD patients carrying ***C9ORF72*** intronic hexanucleotide expansions; cdsRNA isolated from these tissues is comprised of repeat sequences. Together, these findings implicate cdsRNA generated from genomic sequences in neurons as a trigger for sterile, viral-mimetic IFN-I induction and propagated neuronal death within in a neural circuit in the aging nervous system.

## INTRODUCTION

Human neurons can survive for over 100 years. However, the effects of aging (Lopez-Otin et al., 2013) take a toll on the brain and can lead to debilitating adult-onset neurodegenerative diseases such as Amyotrophic Lateral Sclerosis (ALS), Frontal Temporal Dementia (FTD), and Alzheimer’s disease (AD) (Coppede and Migliore, 2015; Suberbielle et al., 2013; Wang et al., 2013). Excessive DNA damage is a feature of aging brains and adult-onset neurodegenerative diseases (Lu et al., 2004; Madabhushi et al., 2014). Aberrant DNA repair can also lead to structural changes in the genome, namely copy number variations, deletions, insertions, and inversions, that are commonly associated with neurological diseases (Chiang et al., 2012; Haeusler et al., 2016; Kovtun et al., 2007; Rovelet-Lecrux et al., 2012; Talkowski et al., 2012). Moreover, germline mutations in DNA repair proteins cause pediatric syndromes with neurodegeneration (Madabhushi et al., 2014). DNA damage and consequential structural rearrangements in the genome are therefore postulated to contribute actively to neurodegenerative processes (Doan et al., 2016; Lodato et al., 2017; Madabhushi et al., 2014).

Neuroinflammation is another hallmark of aging that is also a pathological feature of adult onset neurodegenerative diseases. Gene expression profiling studies of aging murine and human brains reproducibly identified the type-I interferon (IFN-I) antiviral innate immune signaling pathway as the most significantly induced pathway (Baruch et al., 2014; Grabert et al., 2016; Lu et al., 2004). IFN-I pathway activation was also reported in the brains of ALS and AD patients as well as animal models (Hu et al., 2003; O’Rourke et al., 2016; Taylor et al., 2014; Wang et al., 2011). Furthermore, rare genetic disorders collectively referred to as interferonopathies and defined by high systemic levels of IFN-I signaling can present with brain atrophy during childhood (Rice et al., 2014). Viral encephalitides, such as herpes simplex virus (Mancuso et al., 2014) and Zika virus infections (Beckham et al., 2016) have also been associated with activation of IFN-I signaling and massive neurodegeneration. Thus, pathological IFN-I signaling, like excessive DNA damage, is hypothesized to cause neurodegeneration, but how the triggers and degeneration are linked and how this contributes to adult onset neurodegenerative diseases is not known.

Here we report an analysis of mouse lines in which mutant hAPP genes were selectively expressed in mature olfactory sensory neurons (OSNs). Unexpectedly, the integrated transgene was riddled by a complex genomic rearrangement that included inversions, insertions, and deletions. Expression of this rearranged gene results in production of cytoplasmic double stranded RNA (dsRNA), which activates IFN-I antiviral innate immune signaling pathway and causes neuronal death in both transgene-expressing and neighboring cells. The non-cell-autonomous effects of dsRNA expression extend as far as the olfactory bulb of the brain where OSN axons terminate. Comparison of multiple unique hAPP transgenic mice suggests this profound neurodegeneration is independent of hAPP. Consistent with this, we demonstrate that dsRNA derived from non-pathogenic genes is sufficient to induce neurodegeneration *in vivo*. Transfection of a dsRNA mimetic (dsRNAmi) into the cytoplasm of human neurons induces similar IFN-I signaling and cell death. Moreover, we detect cytoplasmic dsRNA and elevated IFN-I signaling in the brains of ALS-FTD patients carrying an intronic hexanucleotide repeat expansion in *C9ORF72* gene. These data indicate that sterile inflammation in neurons initiated by a viral mimetic, cytoplasmic dsRNA, can elicit cell autonomous and non-autonomous cell death.

## RESULTS

### Generation of Neurodegeneration 1 (Nd1) and Nd2 Transgenic Mice

We previously reported that the CORMAP and CORMAC mouse lines, which overexpress human amyloid precursor protein (hAPP) isoforms, in mature OSNs, did not exhibit accelerated death of OSNs (Cao et al., 2012). Simultaneously, we generated two separate hAPP-overexpressing transgenic mouse lines that did exhibit neurodegeneration despite expressing the same alleles, which we describe here. The first mouse line, named <u>N</u>euro<u>d</u>egenerative 1 (Nd1), overexpressed the Swedish isoform of hAPP-K670N/M671L (hAPPsw), which causes familial early-onset AD (Mullan et al., 1992), and the fluorescent marker GFP (Figure 1A). The second mouse line, named Nd2, overexpressed hAPP carrying an M671V mutation and the fluorescent marker protein mCherry (Figure S1A). The synthetic M671V mutation conferred resistance to cleavage by BACE1 (Citron et al., 1995). Thus, Nd1 and Nd2 express distinct isoforms of hAPP that increase and reduce production of the amyloid-forming Aβ peptide relative to wild type hAPP, respectively. Both constructs were placed downstream of a tetracycline responsive element (TRE). Transgene expression was restricted to mature OSNs by crossing Nd1 and Nd2 animals into an OMP-IRES-tTA background (Gogos et al., 2000), which expresses the tetracycline-controlled transcriptional activator protein from the native olfactory marker protein (OMP) locus to create a tet-off system (Figure 1A; Figure S1). This tet-off system allowed us to regulate transgene expression temporally by feeding mice chow containing doxycycline to control for a position effect of transgenic integration.

**Figure 1.**
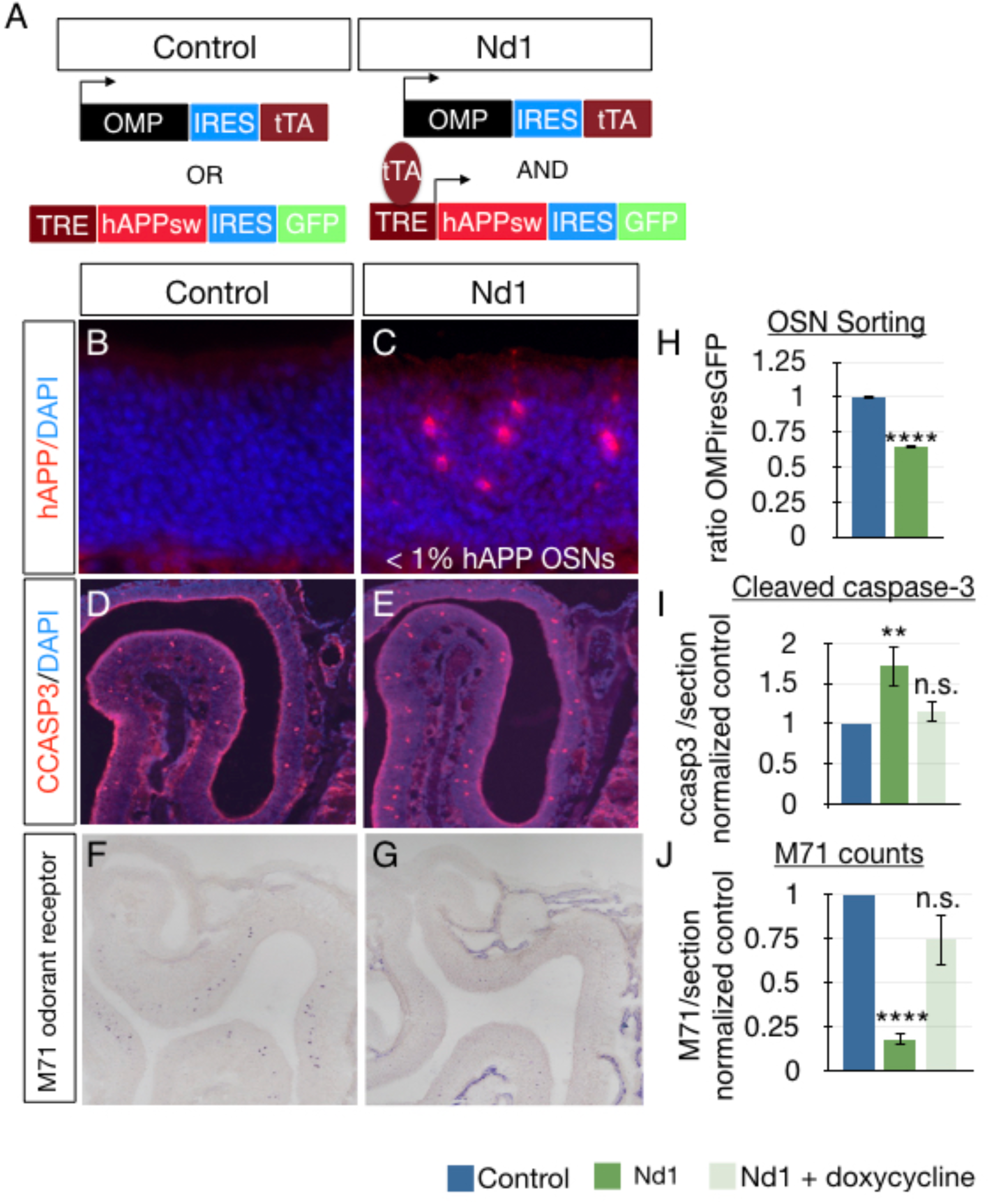
Expression of Nd1 is necessary and sufficient to induce mouse olfactory sensory neuron apoptosis. A) A schematic showing the Nd1 transgenic construct; hAPPsw and GFP are co-expressed with an IRES sequence, which was placed under the transcriptional control of the tetracycline-responsive promoter element (TRE). The tetracycline-controlled transactivator protein is under the control of the endogenous OMP promoter, thereby restricting transgene expression to mature OSNs. We used mice with the TetO transgene alone, or expressing tTA alone littermates as controls. B,C) Coronal sections of mouse olfactory epithelia stained by tyramide amplified immunofluorescence with an antibody specific to hAPP (6E10, red) and the nuclear dye, DAPI (blue), in either littermate control (B) or Nd1 transgenic mice (C). D,E) Coronal sections of mouse olfactory epithelia stained by immunofluorescence with an antibody to cleaved-caspase 3 in littermate control (D) or Nd1 transgenic mice (E). F,G) Coronal sections of mouse olfactory epithelia stained by RNA *in situ* hybridization with a digoxigenin labeled probe for the M71 odorant receptor RNA in littermate control (F), or Nd1 mice (G). H) Percentage of olfactory sensory neurons (all mature neurons labeled with GFP) relative to all cell in the olfactory epithelium quantified by FACS analysis, in Nd1 (green bar) and littermate controls (blue bar). p<0.0001, n=4. I) Quantitation of the number of cleaved-caspase3 positive cells per section of olfactory epithelium in Nd1 normalized to littermate controls (green bar) or for animals fed food containing doxycycline for three months (light green bar). Nd1 p=0.011, n=5; Nd1 + dox p=0.25, n=3. J) Quantitation of the number of M71 positive cells per section of olfactory epithelium in Nd1 normalized to littermate controls (green bar) or for animals fed food containing doxycycline for three months (light green bar). Nd1 p=0.0019, n=5; Nd1 + dox, p= 0.074, n=4.

### Non-cell-autonomous Neurodegeneration in the Mouse Olfactory Neuronal Circuit

Transgene expression in both Nd1 and Nd2 mice caused significant neurodegeneration. Quantification of all mature OSNs in Nd1 and Nd2 mice by fluorescence activated cell sorting (FACS) revealed a striking 40% reduction in the steady-state levels of OSNs at 90 days of age (Figure 1H; Figure S1H) relative to age-matched control littermates. This 40% reduction in steady-state OSN levels persisted in all subsequent ages examined, up to 12 months. At earlier ages, we observed a minimal reduction in OSNs, suggestive of a neurodegenerative process that accrued over time. In P90 animals, OSNs immunostaining for cleaved-caspase 3 (CCASP3), which is a marker of apoptosis, increased ~2-fold relative to littermate controls, respectively (Figure 1; Figure S1). Furthermore, we examined one subclass of OSNs defined by expressing the M71 odorant receptor using RNA *in situ* hybridization. We observed a > 80% and > 50% reduction in OSNs expressing M71 in the Nd1 and Nd2 lines, respectively.

Next, we characterized the expression pattern of hAPP alleles in Nd1 and Nd2 mice using immunofluorescence to detect hAPP. Surprisingly, we observed < 1% of OSNs expressed the Nd1 and Nd2 transgenes (Figures 1B-1C; Figure S1). Quantification using FACS of GFP or mcherry-positive OSNs relative to the total population of OSNs confirmed this estimate (0.83% ± 0.21% SEM, n=4 Nd1 mice, Nd2 0.73% ± 0.04% SEM, n=3 Nd2 mice). The loss of nearly half of OSNs in mice in which transgene expression is limited to <1% of the population indicated a robust non-cell-autonomous neurodegenerative phenotype. Dual labeling of GFP (to identify transgene-expressing cells) and CCASP3 (to identify apoptotic cells) confirmed that the vast majority of apoptotic OSNs in Nd1 did not express the transgene (98.66% ± 0.47%) (Figure S2).

### Complex Genetic Rearrangements, Not hAPP Levels or Position Effect, Correlate with Neurodegeneration

We wondered whether differences in transgene expression levels might cause neurodegeneration in Nd1 and Nd2 but not in animals described in our previous report (Cao et al., 2012). However, we observed no differences in levels of hAPP-K670N/M671L expression between Nd1 and CORMAP mice (Figures S3A–S3C) or in levels of hAPP-M671V between Nd2 and CORMAC (Figures S3D–S3F) or in levels of hAPP-M671V between Nd2 and CORMAC (Figures S3D) mice as judged by immunofluorescence of OSNs using an antibody specific for hAPP (6E10). In addition, no differences in the levels of hAPP RNA were detected by quantitative PCR (qPCR) of transgene positive OSNs isolated by FACS from Nd2 and CORMAC animals (Figures S3G–S3H). Thus, differences in levels of hAPP expression do not explain the loss of neurons in the Nd1 and Nd2 lines relative to CORMAP and CORMAC animals. Moreover, Tg2576 mice, which broadly overexpress hAPPsw and produce very high levels of Aβ peptide in the olfactory epithelium do not exhibit OSN death (Cao et al., 2012). Taken together, these data suggest that Nd1 and Nd2 transgenes trigger neurodegeneration via a shared process that depends neither upon transgene levels nor levels of the amyloid-forming Aβ peptide.

Next we asked whether a position effect caused neurodegeneration in Nd1 or Nd2 transgenic mice. Quantitation of mature OSNs and analysis of CCASP levels revealed no difference between control animals and mice harboring TetO transgenes only, which do not express hAPP due to the lack of the TTA driver. Moreover, in adult Nd1 mice carrying a TTA driver with extensive OSN cell death, the oral administration of doxycycline for three months repopulated OSNs and reduced apoptosis to control levels (Figures 1I-1J). Collectively, these data strongly suggest that transgene transcription is necessary for OSN neurodegeneration in Nd1 and Nd2 animals.

Transgenic constructs can undergo complex genetic rearrangements as a result of chromothripsis during transgene integration, as described in the R6/2 Huntington’s disease (HD) mouse model (Chiang et al., 2012). The rearrangements can include deletions, insertions, and inversions of one or more trangene sequences (Chiang et al., 2012). A similar chromothripsis of human genomes occurs in autism and related neurodevelopmental disorders, suggesting that structural alterations in the genome can alter neuronal function (Chiang et al., 2012; Lohmann et al., 2017; Talkowski et al., 2012). We wondered whether a similar complex genetic rearrangement occurred in Nd1 and Nd2 transgenes leading to a toxic gain-of-function. Accordingly, we performed deep-sequencing on whole genome jumping libraries with insert sizes selected at ~2kb (Hanscom and Talkowski, 2014) using DNA isolated from Nd1 and CORMAP transgenic mice. Sanger sequencing showed that the Nd1 transgene integrated into Chr. 15 and the CORMAP transgene integrated into Chr. 9 (Figure 2, Table SI). However, the structure of the Nd1 transgene revealed a complex genetic rearrangement involving more than two breakpoints with resulting deletions, insertions, and at least one inversion (Figure 2B). In contrast, we did not detect structural alterations of the CORMAP transgene (Figure 2B). The complexity and repetitive nature of transgenes precluded us from determining the complete Sanger sequence.

**Figure 2.**
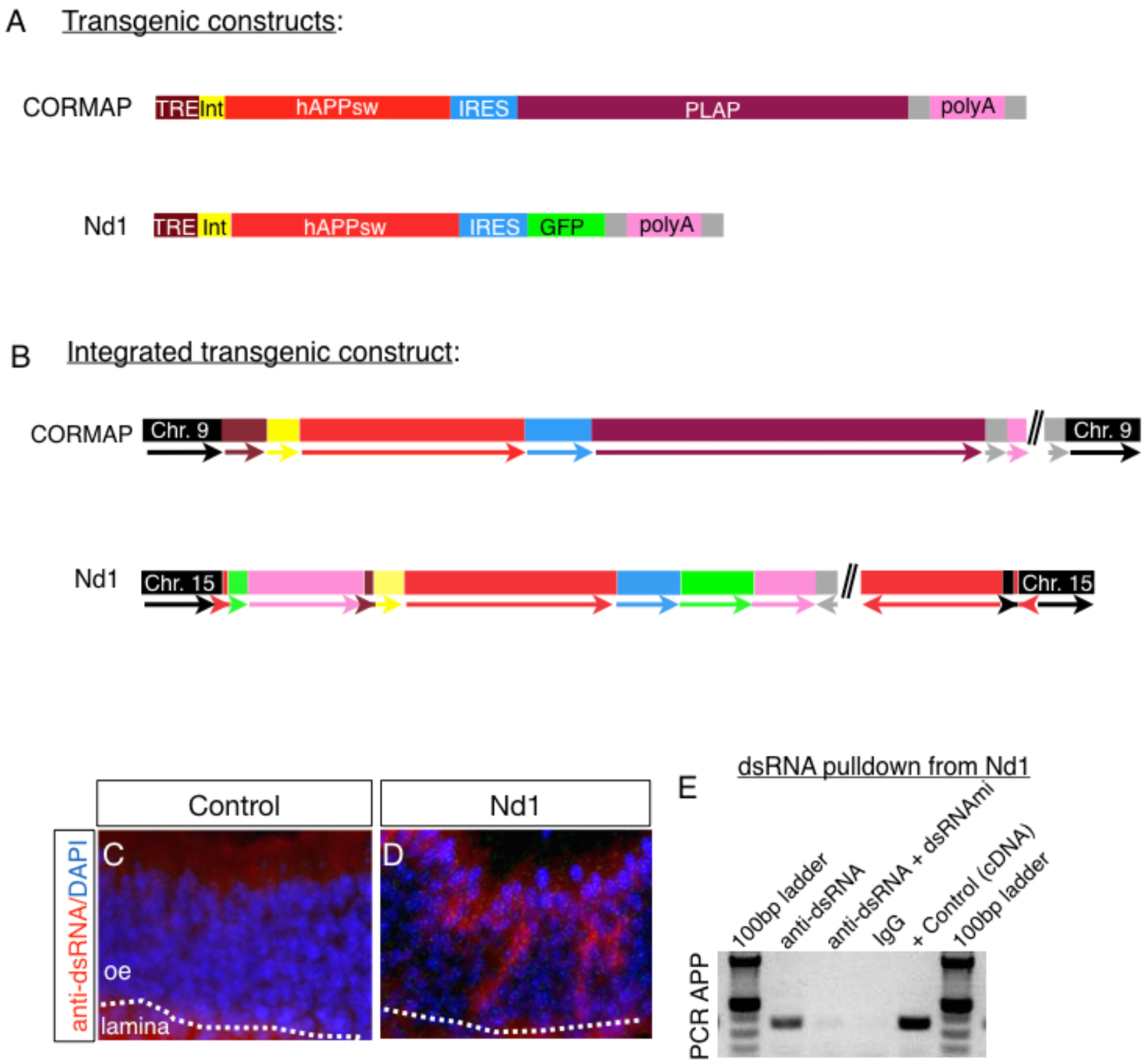
Complex genetic rearrangements in Nd1, but not detected in CORMAP, generate cytoplasmic dsRNA. A) Color coded schematics of cloned transgenic constructs that were injected into mouse eggs for CORMAP and Nd1 transgene sequences, as determined by sanger sequencing. B) Color coded schematics of the structure of the <u>integrated</u> transgenic sequences and their chromosomal locations (flanking black boxes), as determined by whole genome sequencing of jumping libraries and confirmed by sanger sequencing. Double slash (//) represents regions of the transgene where we could not obtain sanger sequence (1 pixel/ nucleotide, then scaled equally to fit). C) Coronal sections of mouse olfactory epithelia stained by tyramide amplified immunofluorescence with an antibody specific to dsRNA (J2). D) PCR amplification of transgenic hAPP from cDNAs generated from RNA isolated by immunoprecipitation with an antibody to dsRNA (J2), but not from immunoprecipitation with isotype and concentration matched IgG2a, or immunoprecipitates from dsRNA (J2) antibodies preincubated with a synthetic dsRNA mimetic.

### Genomically-Encoded Cytoplasmic Double Stranded RNA is Toxic to Neurons *in vivo*

We hypothesized that transcription of inverted DNA sequences might produce intra or intermolecular dsRNA when the antisense strands anneal to the predicted sense strands (Deleidi et al., 2010; Gantier and Williams, 2007; Melton et al., 2003). We performed immunofluorescent imaging, biochemical analysis, and *in situ* hybridization of olfactory epithelia to test this hypothesis. Immunofluorescent staining with an antibody that specifically binds to dsRNA in a sequence-independent fashion revealed that cytoplasmic dsRNA is present in OSNs in Nd1, but not CORMAP, animals (Figures 2C-2D). Similarly, dsRNA-positive OSNs were found in Nd2, but not CORMAC, OSNs (Figures S1F-SG). To confirm that the Nd1 transgene gives rise to dsRNA, we amplified inverted regions (deriving from the hAPP-expressing part of the construct) directly from cDNA libraries generated from dsRNA that had been immuno-precipitated from cytoplasmic extracts prepared from OE tissue using the anti-dsRNA antibody (Bonin et al., 2000). As controls, we showed that inverted DNA could not be amplified when an isotype control antibody was used for immunoprecipitation or when the antibody against dsRNA was pre-incubated with a dsRNA mimetic as a competitor (Figure 2E). Moreover, using RNA *in situ* hybridization to label APP and GFP mRNA in the Nd1 olfactory epithelium, we detected cytoplasmic expression of both sense and anti-sense Nd1 (Figure S4). We found that fewer cells expressed anti-sense GFP and APP relative to sense RNA, suggestive of 1.) a rapid loss of neurons expressing dsRNA above a threshold, 2.) a competitive interaction of endogenous dsRNA with the RNA detection probe, and/or 3.) editing of the dsRNA (Savva et al., 2012). A similar observation has been made in the brains of ALS patients which exhibit substantially fewer neurons expressing antisense ***C9ORF72*** relative to sense ***C9ORF72*** transcripts (as measured using LNA *in situ* probes) (Mizielinska et al., 2013). Taken together these results demonstrate that the transgene in Nd1 but not CORMAP mice is rearranged and that inversions resulting from transgene rearrangements generate dsRNA in the cytoplasm of Nd1 OSNs. Interestingly, dsRNA may explain the restricted expression of Nd1 and Nd2 transgene to only 1% of neurons (Calero-Nieto et al., 2010).

Next, we wondered whether cytoplasmic dsRNA was sufficient to induce neurodegeneration. We developed an alternative strategy to express both sense and antisense GFP in mature OSNs *in vivo* (Figure 3A). We hypothesized that transcription of antisense-GFP would lead to formation of dsRNA species in select OSNs that stably transcribe sense-GFP. Accordingly, mice expressing GFP exclusively in mature OSNs (OMP-IRES-GFP mice (Shykind et al., 2004)) were infected with an AAV9-flex-GFP virus (Cardin et al., 2009), which constitutively expresses antisense GFP in the absence of cre (Figure 3A). As negative controls, we instilled AAV9-GFP at the same viral titer into OMP-ires-GFP mice, and we instilled AA9-flexGFP into mice expressing tTA from (but not GFP) from the OMP locus. Three to four weeks after viral infection, we observed cytoplasmic dsRNA via immunofluorescence in AAV9-flex-GFP-infected animals, but not in negative controls (Figures 3B-3C). We confirmed the presence of antisense GFP using RNA *in situ* hybridization. We observed a ~30% reduction in mature OSNs in mice expressing cytoplasmic dsRNA relative to controls (Figures 3D-3F) using RNA *in situ* hybridization with a probe against OMP to label mature OSNs. These data are consistent with previous studies showing that intranasal instillation with polyinosinic-polycytidylic acid, a synthetic dsRNA mimetic (dsRNAmi), triggers degeneration of both OSNs in mice and degeneration of cortical neurons in the rat (Kanaya et al., 2014; Melton et al., 2003).

**Figure 3.**
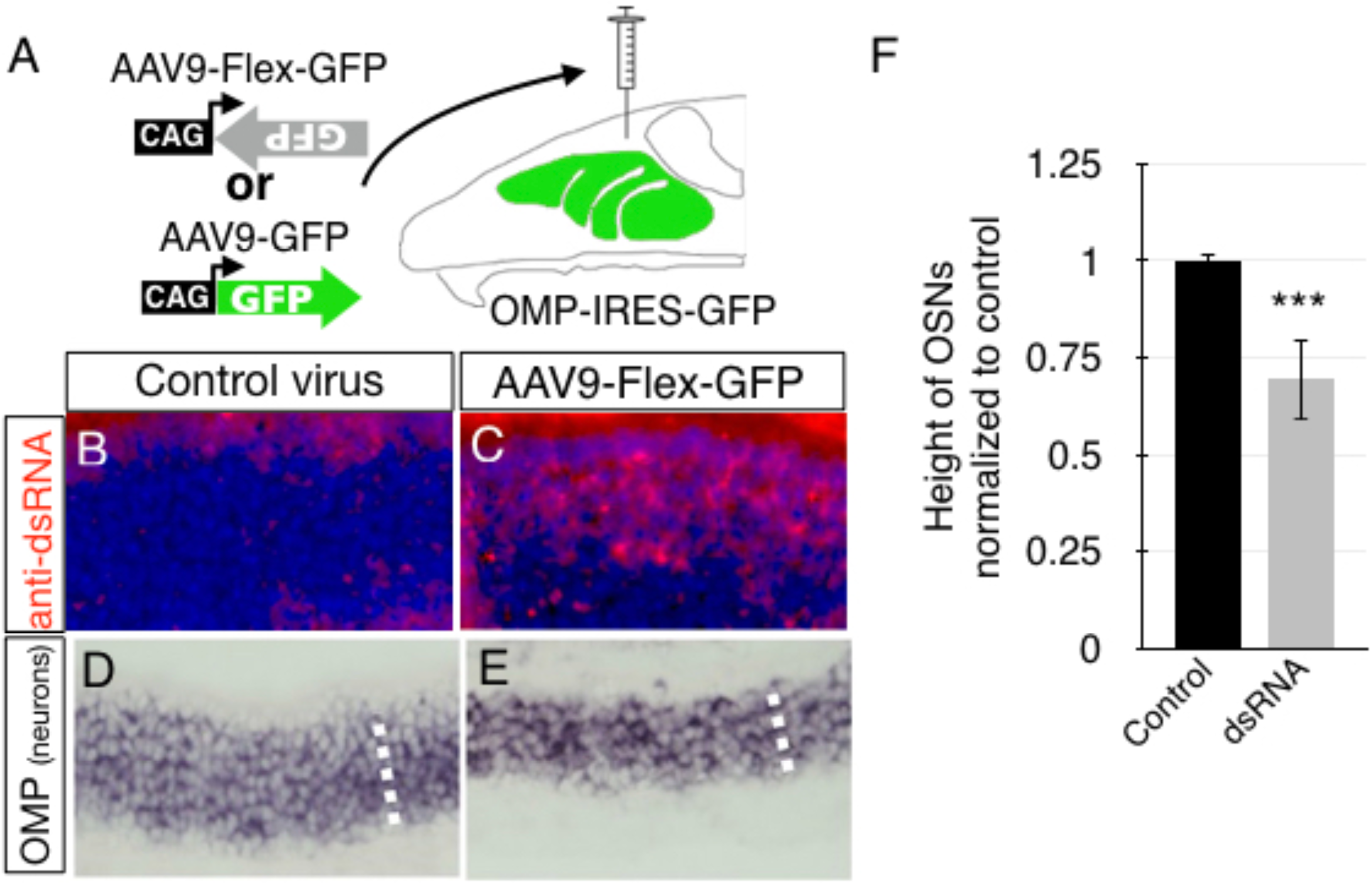
DsRNA is sufficient to induce death of olfactory sensory neurons. A) schematic showing mice expressing GFP in mature OSNs were instilled with AAV9 viral constructs for the expression of with either sense or antisense GFP. B,C) Coronal sections of mouse olfactory epithelia stained by tyramide amplified immunofluorescence with an antibody specific to dsRNA (J2) in control (B) or mice instilled with antisense AAV9 (C). D,E) Coronal sections of mouse olfactory epithelia stained by RNA *in situ* hybridization with a digoxigenin labeled probe for olfactory marker protein (OMP) in control (D) or mice instilled with antisense AAV9 (E). F) Quantitation of height of olfactory epithelia positive for OMP staining normalized to controls, representing loss of OSNs ((p<0.0001, n=4 pairs of animals).

### Genomically-Encoded dsRNA in Neurons induces the Type I Interferon Pathway *in vivo*

Neuroinflammation is a pathogenic hallmark of neurodegenerative disease (Ransohoff, 2016). We wondered whether dsRNA upregulates innate immune signaling pathways in OSNs of Nd1 animals (Deleidi et al., 2010; Gantier and Williams, 2007; Melton et al., 2003). Accordingly, we carried out RNA-seq experiments comparing the transcriptomes of purified mature OSNs from Nd1 and littermate control mice (Table S2). Briefly, to identify changes in gene expression in an unbiased manner we bred Nd1 animals with OMP-IRES-GFP-expressing mice (express GFP in 100% of OSNs) (Shykind et al., 2004) and sorted GFP+ cells by FACS to isolate mature OSNs to 98% homogeneity (Figure 4A). Following RNA-seq, we used Ingenuity Pathway Analysis (IPA) software to identify upregulated molecular pathways induced by transcriptional changes that were two-fold or greater in Nd1 relative to littermate control samples. Strikingly, the IFN-I antiviral signaling pathway was the most highly upregulated pathway between Nd1 and control (Figure 4B; Table S3). We subsequently confirmed the increased expression of three highly upregulated IFN-I induced genes in the Nd1 olfactory epithelium, but not CORMAP, including signal-transducing activators of transcription 1 (*Stat1*), 2’-5’ oligoadenylate synthetase-like 2 (*Oasl2*), and interferon induced protein 44 (*Ifi44*), by RNA *in situ* hybridization (Figure 4). IFN pathway activation was confirmed by Western blotting, which revealed a dramatic ten-fold increase in the level of STAT1 phosphorylation on an activating site (pSTAT1 – Tyr701) in extracts derived from Nd1 olfactory epithelia relative to littermate control mice (Figure 4M). Consistent with the notion that IFN-I antiviral activity was not a consequence of hAPPsw expression, *Stat1*, *Oasl2*, and *Ifi44* were also highly expressed in OSNs of Nd2 mice, but not CORMAC or CORMAP mice (Figures S5). Given that IFN-I signaling can be stimulated by either IFNα, IFNβ, and that there is overlap with IFNγ-mediated signaling, we examined our RNA-seq data for cytokine expression. IFNα4 and IFNα12, but not IFNγ or IFNβ, were elevated in Nd1 neurons (Table S4), consistent with IFN-I signaling resulting from the stimulation of antiviral interferons by nucleic acids.

**Figure 4.**
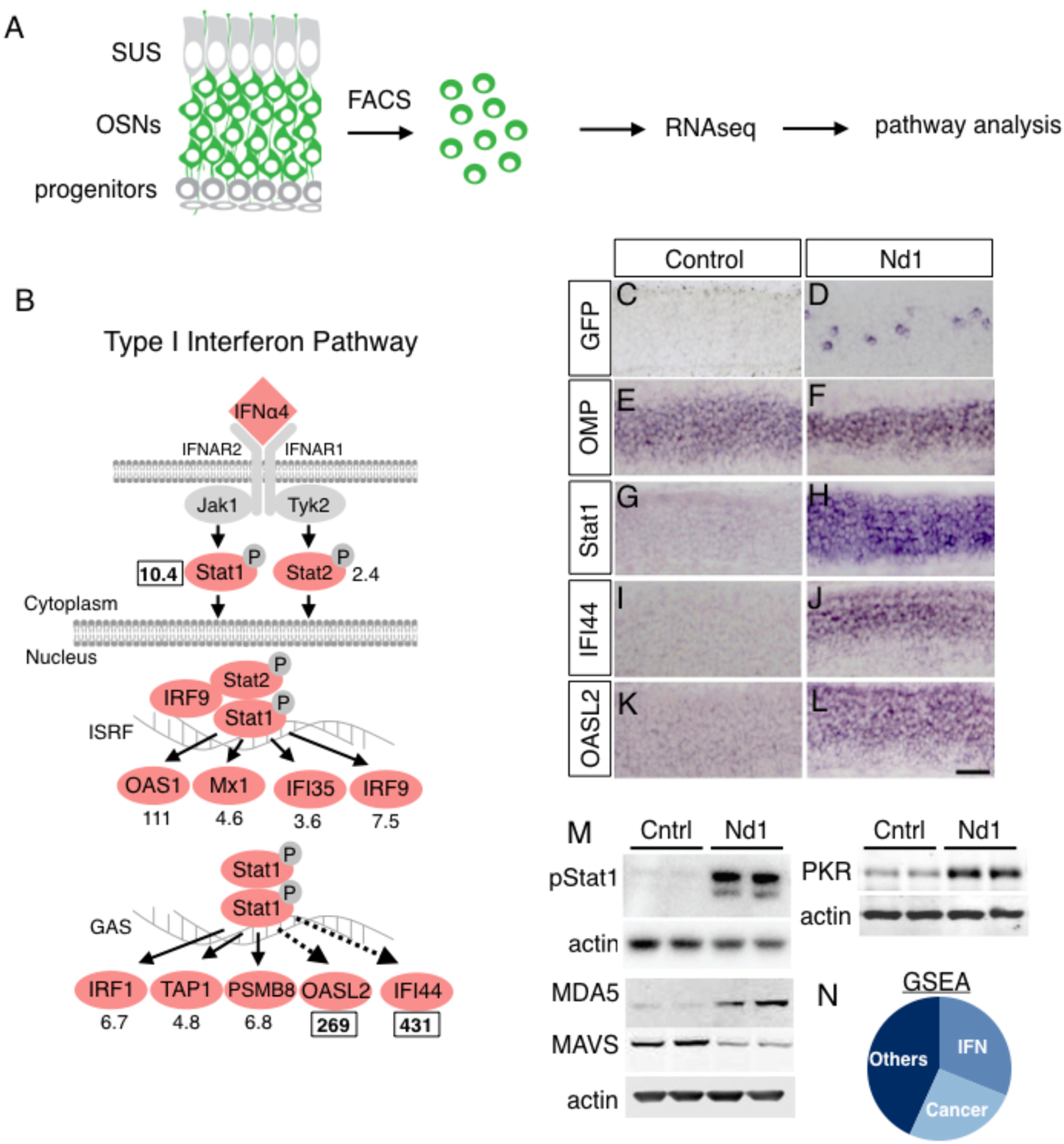
Type I interferon signaling is highly unregulated in Nd1 relative to littermate controls. A) Schematic showing mature OSNs expressing GFP were isolated by fluorescence-activated cell sorting. RNA from sorted neurons was processed for RNA-seq and analyzed. B) Schematic modified from Ingenuity Pathway Analysis software, showing the upregulation type I interferon signaling pathway in Nd1 (p = 6.2x10^-6^). Genes labeled in red are up-regulated by indicated fold values. Genes with boxed fold values were used to confirm the RNA-seq by RNA *in situ* hybridization (G-L) or western blotting (M): C,D) GFP expression showing transgene expression (~1% of cells). B, C) OMP was used as a positive control to label mature OSNs. The expression of Stat1 (G, H), IFI44 (I, J), and OASL2 (K, L) are all unregulated in Nd1 relative to controls by RNA. M) Western blot showing elevated phosphorylation of Stat1, MDA5, PKR, and reduced small isoform of MAVS in Nd1 relative to controls. N) Pie chart of the distribution of pathways significantly upregulated by GSEA analysis.

### Propagation of Neurodegeneration in the Brain in Nd1 and Nd2 mice

Transgene expression in the Nd1 line is detectable in a very small fraction of OSNs (< 1%), but the distribution of apoptosis (40% reduction in steady-state levels of OSNs; Figure 1 and Figure S2) and IFN-I signaling (Figures 4D, H, J, L) in OSNs is much greater, indicating a non-cell autonomous process in the OE. We wondered whether neurodegeneration and IFN-I signaling also propagated in a non-cell-autonomous manner to the olfactory bulb of the brain, where OSN axons terminate (Figure S6). Accordingly, we assayed multiple neuronal subtypes in the olfactory bulb by RNA *in situ* hybridization for *Oasl2* as a marker of the IFN signaling pathway in Nd1 and Nd2 animals. Strikingly, we observed elevated *Oasl2* levels in several cell types in both Nd1 and Nd2 (Figures 5D-5G; Figures S7A–S7B), including: 1) periglomerular (Pg) interneurons surrounding OSNs axon termini, 2) mitral cells (Mi) that are the postsynaptic partner of OSNs, and 3) granule interneurons (Gr) which lie deep within the olfactory bulb and do not directly synapse with OSNs. Importantly, we confirmed that the transgene is not expressed in these cell types (Figures 5A-5C; Figures S6B–S6D).

**Figure 5.**
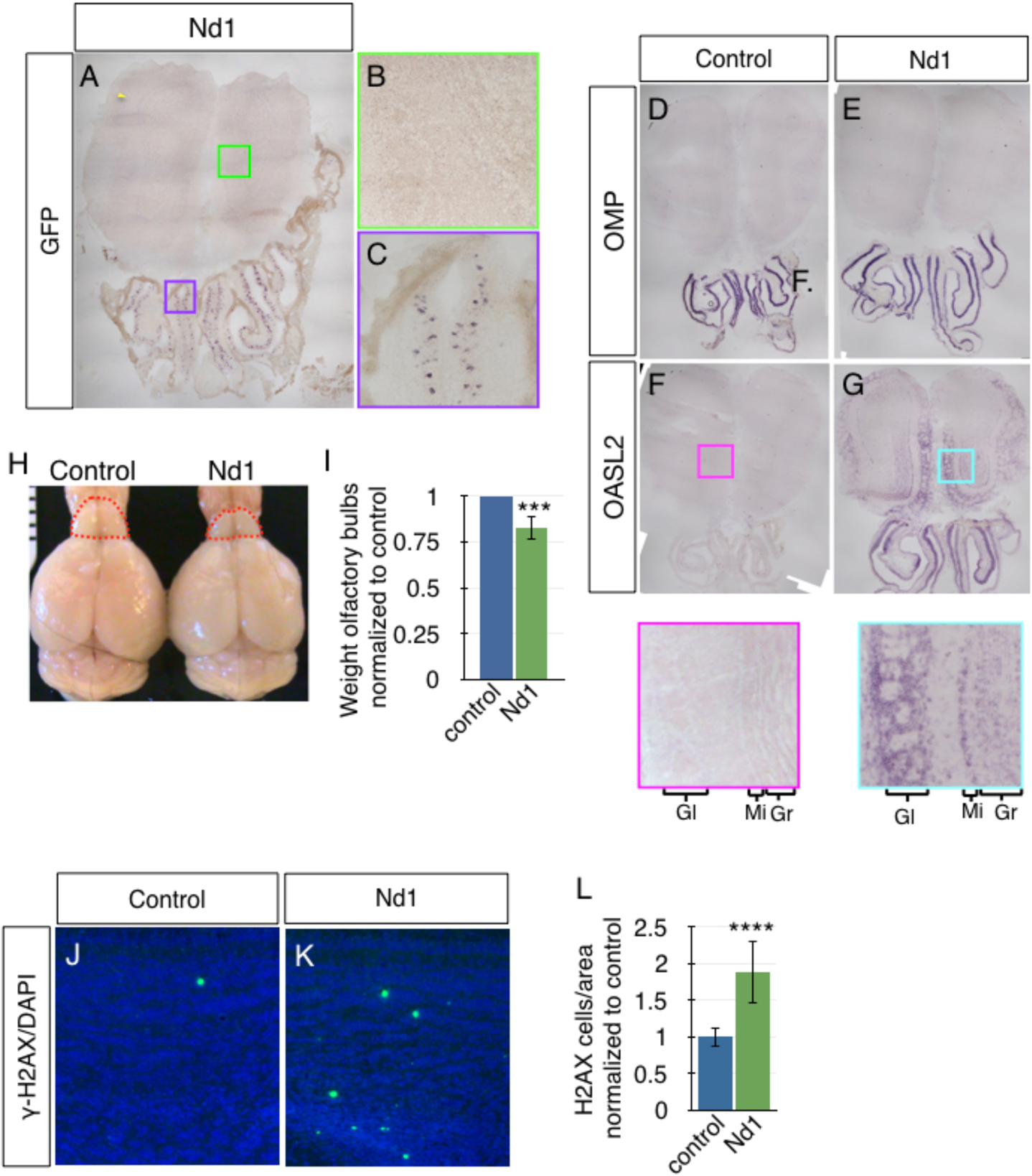
Propagation of IFN-I signaling and neurodegeneration in Nd1. A-C) Coronal sections of mouse olfactory epithelia stained by RNA *in situ* hybridization with a digoxigenin labeled probe for GFP (transgene). Transgene expression is restricted to the cells bodies of OSNs in the olfactory epithelium (A,C), but not cells in the olfactory bulb (A,B). Coronal sections of mouse olfactory epithelia stained by RNA *in situ* hybridization with a digoxigenin labeled probe for: D,E) OMP, a positive control for OLFACTORY SENSORY NEURON expression in control (D) and Nd1(E); F,G) OASL2, as a marker of IFN-I activity- note that interferon expression propagates from the olfactory epithelium to all layers in the olfactory bulb (insets). H) Olfactory bulbs were smaller in Nd1 relative to controls and olfactory bulbs were weighed (I) (p= 0.001, n=4). (J,I) Coronal sections of mouse olfactory epithelia stained by immunofluorescence with an antibody for γH2AX in control (J) and Nd1 (I). K) Quantitation of cells with pan-nuclear γH2AX staining per coronal section of olfactory bulb normalized to the area of the olfactory bulb in Nd1 relative to littermate control mice. p < 0.0001, n=3 mice, at lease 6 olfactory bulb sections per mouse.

We next assessed olfactory bulbs from Nd1 mice for frank neurodegeneration. The olfactory bulbs of Nd1 mice were approximately 20% smaller by weight compared to littermate controls (Figures 5H-5I). CCASP3 is present at high levels in the axons of OSNs in Nd1 mice, which obscures CCASP3 in cells of the olfactory bulb. To circumvent this issue, we used immunofluorescent staining for γH2AX, which marks large chromatin domains surrounding damaged DNA (Lu et al., 2006). We observed an 80% increase in neurons exhibiting pan-nuclear γH2AX staining in Nd1 bulbs, consistent with apoptosis (Figures 5J-5L) (Lu et al., 2006). Similarly, we observed propagated IFN-I signaling into the olfactory bulb and a reduction in olfactory bulb size in Nd2 mice (Figures S7C–S7D). Taken together, these data show that IFN-I signaling and neurodegeneration is propagated from the nose to the brain, presumably via OSN axons.

MDA5, RIG-I and PKR are cytosolic pattern recognition receptors (PRRs) that bind endogenous cytoplasmic dsRNAs (Nallagatla et al., 2011; Wu et al., 2013). Activation of these PRRs by dsRNA activates a positive feedback loop that enhances expression of the receptor genes themselves (Takeuchi and Akira, 2010). By Western blot (Figure 4M) we found that PKR and MDA5 protein levels were elevated in Nd1 animals relative to littermate controls. In our RNA-seq data (Table S2), both MDA5 and RIG-I are induced, which are two dsRNA PRRs that mediate signaling through the mitochondrial anti-viral signaling protein (MAVS). These activated dsRNA PRRs trigger the aggregation and degradation of the large isoform of MAVS (Brubaker et al., 2014; Castanier et al., 2012; Wu et al., 2013). When we measured the levels of MAVS by Western blotting of whole cell extracts from Nd1 olfactory epithelia, we observed an approximately 70% reduction in the levels of the MAVS large isoform relative the smaller isoform, compared to littermate control mice (Figure 4M). From these data we conclude that dsRNA generated by transgene expression in Nd1 mice activates PRRs, such as MDA5 or RIG-I, degrades MAVS, and induces IFN-I signaling in OSNs.

### Cytoplasmic DsRNA Induces Type I interferon Signaling and Death in a Dose-dependent Manner in Cultured Human Neurons

To determine if human neurons also induce IFN-I signaling in response to cytoplasmic dsRNA, we transfected dsRNAmi (polyinosinic:polycytidylic acid) into the cytoplasm of differentiated human ReNcell VM neurons (Kim et al., 2015) derived from normal neuroprogenitors that were transformed by v-myc. We confirmed the cytoplasmic localization of the dsRNAmi using a Rhodamine tagged dsRNAmi (Figure S8). At 24 hours after dsRNAmi transfection, we observed marked induction of several IFN-I induced proteins, including MDA5, PKR, and pStat1 (Figure 6A). At 48 hours post transfection, we found a dose-dependent death of neurons, consistent with the loss of OSNs in Nd1 mice (Figure 6B; Figure S8). Similarly, transfection of dsRNAmi into ReNcell CX neurons, a second human neuronal line derived from normal neuroprogenitor cells that differentiated into glutamatergic cortical-like neurons, also induced neuronal death after 48 hours (Figure S9). Taken together, these data suggest that dsRNA expressed in human neurons induces a robust IFN-I response leading to apoptosis.

**Figure 6.**
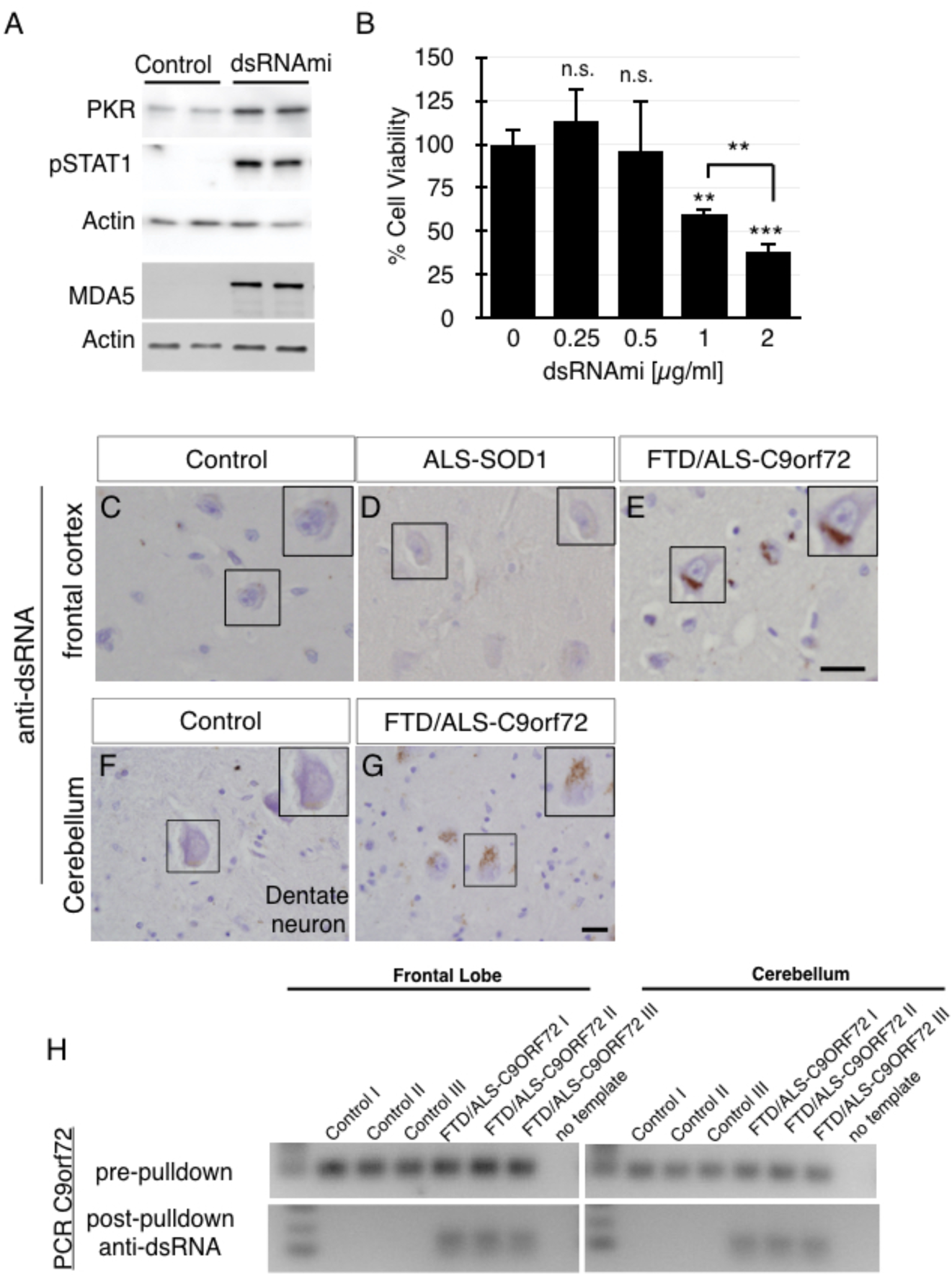
DsRNA is sufficient to induce neuronal cell death in differentiated human neurons and is present in the cerebelli of ALS/FTD patients with the ***C9ORF72*** mutation. A) Western blots showing the induction of IFN-I signaling in differentiated human neurons (ReNcell VM) 24 hours post transfection with a dsRNAmi. B) Quantitation of neuronal viability using the CellTiter-Glo reagent to measure ATP levels 48 hours post transfection with a dsRNAmi at 0.25 (p=0.4807, n=4), 0.5 (p=0.891, n=4), 1 (p=0.0031, n=4), and 2 μg/mL (p=0.0005, n=4). Note that transfection of 2 μg/mL dsRNAmi causes significantly more neuronal death that 1 μg/mL (p=0.01, n=4). Immunohistochemistry with antibody specific to dsRNA (J2) on fixed thin sections of frontal cortex from: (C) normal control brains, (D) ALS patient with an ***SOD1*** mutation, and (E) an ALS/FTD patient with a *C9ORF72* mutation (E) (scale bar = 25 micron). Anti-dsRNA immunostaining on sections from cerebellum of (F) control and (G) ALS/FTD patient with ***C9ORF72*** mutation. (scale bar = 10 micron). Insets digitally magnified 3 times. H) PCR amplification of cDNA with primers distal to the ***C9ORF72*** expansion show a product made from RNA isolated by immunoprecipitation with an antibody specific for dsRNA from frontal cortex and cerebellum from ALS/FTD- patients with the ***C9ORF72*** mutation, but not normal brains.

### Cytoplasmic DsRNA and Type I interferon Signaling Are Present in Neurons of ALS/FTD-C9ORF72

Having uncovered that cytoplasmic dsRNA can robustly induce IFN-I signaling in neurons and their accelerated death in both mouse OSNs and human cultured neurons, we sought to identify relevance for this mechanism in neurodegenerative disease. The presence of both sense and antisense RNA foci have reproducibly been observed in the nucleus and cytoplasm of neurons from ALS patients carrying an intronic hexanucleotide repeat expansion in the ***C9ORF72*** gene (ALS/FTD-C9ORF72) (Gendron et al., 2013; Lagier-Tourenne et al., 2013; Mahoney et al., 2012b). We hypothesized that transcription of both strands of the intronic hexanucleotide repeat expansion might generate cytoplasmic dsRNA. Using an antibody against dsRNA, we observed elevated levels of dsRNA in the cytoplasm of large cells, presumably neurons, in the frontal cortex and in large cells, presumably neurons, of the cerebellar dentate nuclei in ALS/FTD-***C9ORF72*** patients relative to control patients (Figures 6C-6G). Notably, cdsRNA was not detectable in the molecular, Purkinje, or granule layers of the cerebellum (Figures S10A), where dipeptide repeats (Ash et al.; Gendron et al., 2015; Mori et al., 2013a; Mori et al., 2013b; Zu et al., 2013) and p62 inclusions (Mahoney et al., 2012a) have been observed. We confirmed the specificity of dsRNA immunostaining by pre-incubating the anti-dsRNA antibody with a dsRNAmi competitor prior to staining of tissue sections (Figures S11A). We also confirmed the specificity for dsRNA relative to dsDNA by dot blotting (Figure S11B). Brains from ALS/FTD-C9ORF72 patients contain cytoplasmic inclusions containing the TAR DNA-binding protein 43 (TDP-43) in neurons, which have been associated with increased levels of dsRNA, in part due to derepression of endogenous repeat sequences in the genome. We did not detect an enrichment of dsRNA in the frontal cortex of two ALS patients with a pathogenic SOD1 mutation, a variant of ALS that does not contain TDP-43 inclusions (Figures 6C-6D).

Subsequently, to determine if pathogenic expansions of ***C9ORF72*** are a source of dsRNA we performed dsRNA-immunoprecipitation (dsRIP) and converted dsRIP isolated dsRNA into cDNA for PCR analysis. Using primers distal to the hexanucleotide repeat in ***C9ORF72***, we amplified the hexanucleotide repeat from dsRIP performed on both the cerebellum and frontal cortex (Figure 6H). Immunoprecipitation with an isotype control antibody did not produce detectable PCR product. Taken together, these results indicate that transcription of the hexanucleotide repeat generates dsRNA in the cerebellum and frontal cortex of ALS/FTD-C9ORF72 patients, which is visible in the cytoplasm.

We reasoned that the cytoplasmic dsRNA that we identified in the brains of C9ORF72 mutation carriers would result in the activation of IFN-I signaling. Therefore, we performed an unbiased analysis of previously published RNA-seq reads from ALS-C9ORF72 cerebelli (GSE67196; Prudencio et al., 2015). In our analysis, we identified 110 genes with a two-fold or greater change in expression relative to control patients (Prudencio et al., 2015) (Figure 7A). Only 4 genes were down-regulated, one of which was ***C9ORF72***. Using Enrichr, a web-based pathway analysis software (Chen et al., 2013; Kuleshov et al., 2016), to analyze this differential gene list, the most significantly upregulated “GO Molecular Function” pathway was “double-stranded RNA binding” (Figure 7B). Furthermore, interferon signaling (both interferon gamma and alpha, which overlap) were the most significantly upregulated ligand signatures in the LINCS L1000 datasets (Figure 7C; Tables S5-S6). Furthermore, published analysis of RNA-seq from frontal lobes of the same patients suggest elevated Interferon-alpha cytokines in ALS relative to controls (Prudencio et al., 2015). Collectively, these data link the presence of cytoplasmic dsRNA with elevated IFN signaling in the cerebellum of patients with pathogenic C9ORF72 expansions.

**Figure 7.**
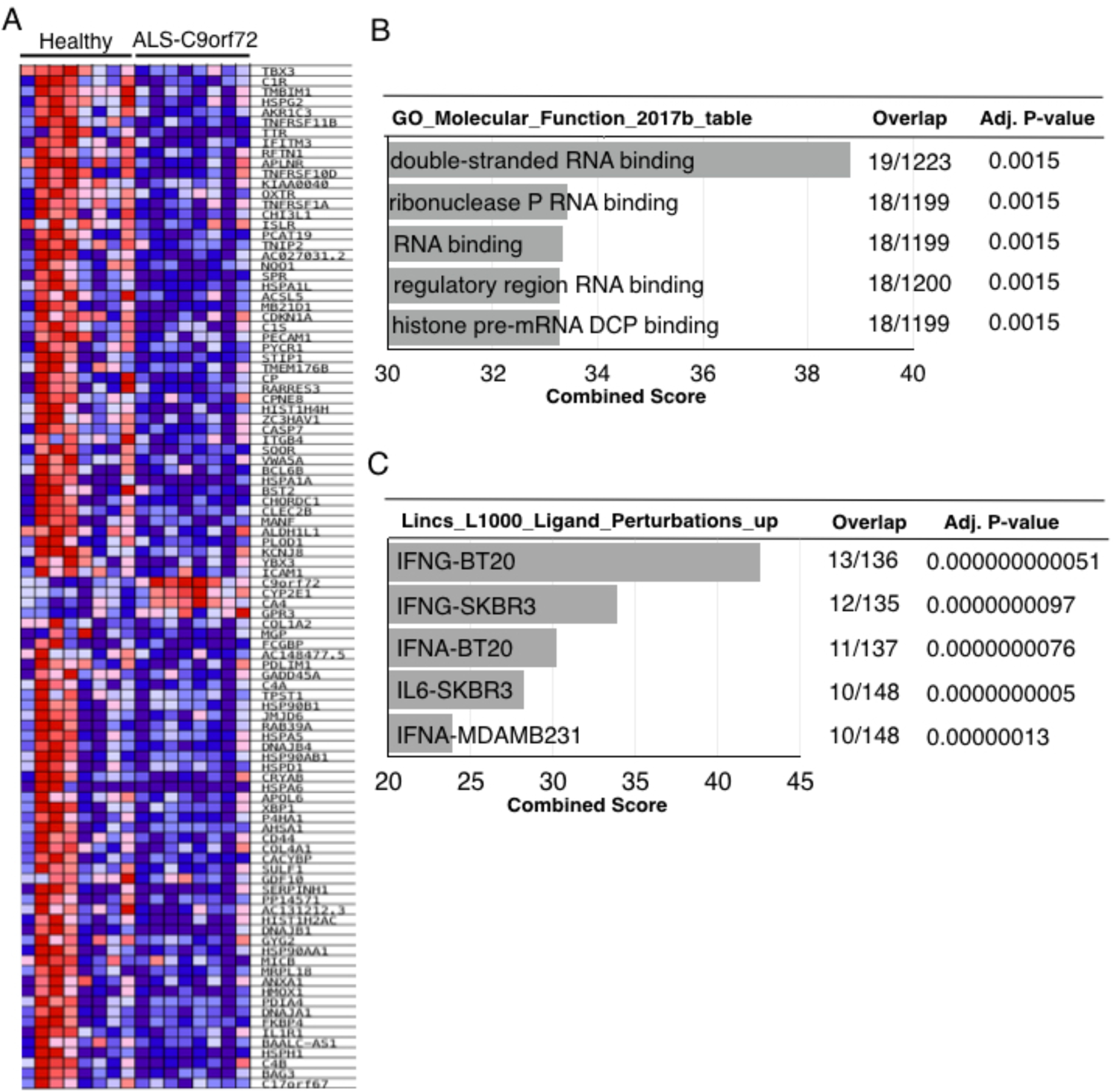
Analysis of gene expression from the cerebellum shows evidence of dsRNA mediated biology in ALS patients with the ***C9ORF72*** mutation. A) A heatmap showing all of the genes that are differentially expressed at 2-fold or greater in ALS patients with the ***C9ORF72*** mutation relative to normal controls (n=8 for each). B) Analysis of the differentially expressed genes in (A); (B) dsRNA binding is the most represented GO Molecular Function, and (C) Both gamma and alpha interferon signaling, which overlap, are the most significantly increased pathways represented in the signatures in upregulated genes in the LINCS L1000 Ligand Perturbations database. We used the publicly available GSE67196 RNA-seq expression data for our analysis.

## DISCUSSION

The severity of synaptic loss and of neuronal death correlate with the onset and progression of symptoms in neurodegenerative disease (Masliah et al., 1994). The transition from stochastic loss of a subset of neurons to a propagated neurodegeneration is an important temporal component in the neurodegenerative process. Once initiated, propagated neurodegeneration increases the likelihood that neighboring neurons or their synaptic partners will succumb to the disease progression. This leads to the systemic destruction of a neural system. We report here that cdsRNA produced from an inverted repeat embedded within in a transgene results in the sterile induction of IFN-I signaling in murine olfactory neurons. Activation of the IFN-I pathway in a small subset (1-3%) of olfactory sensory neurons is associated with the death of not only of these cells, but also of neighboring olfactory neurons and neurons more than two synapses downstream in the brain. The sequences comprising the dsRNA element do not appear to be critical because creation of dsRNA by production of sense and antisense GFP is sufficient to drive Type I IFN signaling and neurodegeneration. However, the amount of dsRNA does appear to be important as the transfection of a dsRNA mimetic into the cytoplasm of human neurons induces dose-dependent type I interferon signaling and innate immune mediated-neuronal death. Finally, we demonstrate cytoplasmic dsRNA accumulation and markers of elevated IFN-I signaling in the brains of ALS/FTD patients. Isolated dsRNA from the frontal cortex and cerebellum of ALS/FTD brains with the expansion of a hexanucleotide repeat of ***C9ORF72***. ***C9ORF72*** is the most common genomic lesion known to produce both sense and antisense RNA, which encodes RNA generated from the adjacent region of the expanded hexanucleotide allele. Collectively, these results provide a new framework for a sterile, viral-mimetic of neurodegeneration triggered by an established pathogenic species, cytoplasmic dsRNA.

### Neuronal Toxicity Mediated by Expansion of *C9ORF72* Hexanucleotide Repeats

Three complementary hypotheses have been proposed to explain how the abnormal expansion of the hexanucleotide repeat of ***C9ORF72*** may be toxic to neurons: 1. Translation of dipeptide repeats via RAN translation from both sense and anti-sense strands (Ash et al., 2013; Gendron et al., 2015; Lagier-Tourenne et al., 2013; Mori et al., 2013a; Zu et al., 2013) to generate abnormal peptide that disrupt essential cellular functions including nuclear-cytoplasmic transport (Hautbergue et al., 2017; Zhang et al., 2015) and exacerbate DNA damage (Lopez-Gonzalez et al., 2016); 2. Sequestration of nuclear RNA binding proteins by nuclear RNA foci comprised of pure sense or antisense strands leading to altered RNA splicing and/or nuclear export (Cooper-Knock et al., 2012; Haeusler et al., 2014; Lee et al., 2013; Mackenzie et al., 2013). 3. Loss of function of C9ORF72 which leads to activated innate immune signaling, but does not cause neurodegeneration in (Atanasio et al., 2016; Burberry et al., 2016; Koppers et al., 2015). Our findings nominate a fourth, complementary, mechanism of cytoplasmic dsRNA-mediated neurotoxicity, that is, activation of pattern recognition receptors by this established PAMP to provoke a sterile, viral-mimetic type I interferon response. Importantly, these four mechanisms may act in combination to promote propagated neuronal lethality. For instance, increased DNA damage and splicing defects mediated by dipeptides may increase the production of dsRNA moieties (Hautbergue et al., 2017; Zhang et al., 2015), and the cytoplasmic accumulation of dsRNA species might be accelerated by dysregulated nuclear cytoplasmic transport or transient nuclear envelope breakdown (Kim and Taylor, 2017). The stability of dsRNA, relative to single strand RNA in our mouse model, as well as recent studies implicating viral dsRNA as a mediator of propagated interferon signaling (Nguyen et al., 2017) and stress granule formation (Yoo et al., 2014), suggest that cytoplasmic dsRNA may be a species responsible for propagation to non-dsRNA producing neurons. However, secreted interferon and prion-like low complexity domain conformations of proteins in the IFN-I signaling cascade (Cai et al., 2014; Hou et al., 2011) are also possible mediators.

### IFN Activation in Neurodegenerative Disease

Activation of IFN-I signaling is observed in the nervous system of patients with ALS, Alzheimer’s, Parkinson’s, Huntington’s and other neurodegenerative diseases and may precede the clinical onset of these diseases (Fritz-French and Tyor, 2012; Hu et al., 2003; Peel, 2004; Stobart et al., 2007). Consistent with the notion that IFN-I signaling induces neurodegeneration, acute high-dose treatment of patients with cancer or viral infections with interferon-α can cause neurological abnormalities, including cognitive dysfunction (Lieb et al., 2006; Meyers et al., 1991; Poutiainen et al., 1994; Raison et al., 2005) and anosomia (Cocquyt and Van Belle, 1994; Maruyama et al., 1998). Moreover, mutations in ***MDA5*** that enhance type I IFN signaling cause Aicardi-Goutieres disease (Rice et al., 2014), which presents with profound neurodegeneration in childhood and interferonopathies associated with neuronal dysfunction (McGlasson et al., 2015). However, despite mounting evidence that interferonopathies induce neuronal dysfunction, the source of pathological IFN-I activity in neural tissue remains elusive.

In human brain tissue, we demonstrate that cytoplasmic dsRNA is comprised, in part, of RNA expressed from the allele of the massively expanded hexanucleotide repeat region of ***C9ORF72***. Several additional sources of cytoplasmic dsRNA exist. First, reactivation of retroviral elements within the genome (HERVs) have the potential to form dsRNA, and have been identified in ALS patients (Douville et al., 2011; Li et al., 2015). Second, knock-down of TDP43, a nuclear protein that is frequently mislocalized to the cytoplasm in ALS, FTD, and a significant proportion of people with AD patients, was shown to promote dsRNA formation in *C. elegans*, and two human neuronal cell lines (Saldi et al., 2014). Moreover, mouse brains depleted of TDP-43 show IFN-I activation (Polymenidou et al., 2011), potentially by de-repression of transposable elements (Li et al., 2012). Third, reactivation of viral infections such as Herpes Simplex Virus 1 and Human Herpes Virus 6, which pass through a dsRNA intermediate and induce neurodegeneration that correlates with ALS, AD, and PD pathology, (Weber et al., 2006) may activate IFN-I signaling (Deleidi and Isacson, 2012; Itzhaki, 2014; Kuhlmann et al., 2010; Li et al., 2015). Future sequencing studies that isolate cytoplasmic dsRNA from human brain tissue in these myriad cases will further illuminate the identity and diversity of dsRNA sources.

### Neuroni<u>n</u>flammation as a driver of Neuroinflammation

A recent report demonstrated that peripheral IFN-I signaling in a lupus mouse model also lead to synaptic loss and neurodegeneration, presumably through activation of microglia within the brain (Bialas et al., 2017). In our mouse models, fortuitous genomic structural variants in Nd1 and Nd2 mice, an artifact of engineering transgenic mice, may model complex genomic rearrangements that arise in neurons and precede neurodegenerative disease (Li et al., 2016; Madabhushi et al., 2014). Moreover, the genetic inversions in Nd1 are markedly similar to complex genetic rearrangements found in the genomes of children with neurodevelopmental disorders (Chiang et al., 2012; Talkowski et al., 2014; Talkowski et al., 2012). In the Nd1 and Nd2 mouse models, genomically-encoded dsRNAs expressed exclusively in a subset of OSNs activate IFN-I pathways in a broad population of neurons, which then can trigger activated astrocytes and microglia. We demonstrate that cultured human neurons are also sufficient to mount a robust IFN-I response, a process that we refer to as neuro<u>n</u>inflammation, which is one type of neuroinflammation.

Together, this new framework of genomically-encoded cytoplasmic dsRNA mediated neurodegeneration implicates neurons as a potential activator of the neuroinflammatory cascades observed in ALS/FTD and other neurodegenerative diseases, and possibly Alzheimer’s disease. Moreover, genomic instability, a well-documented feature of aging and accelerated in neurodegenerative diseases (Lodato et al., 2017), may trigger this sterile viral-mimetic dsRNA pathologic process. From a translational perspective, arrest of the innate immune response triggered by accumulated dsRNA, which we speculate will comprise a significant proportion of sporadic ALS patients and minority of AD patients that have cytoplasmic TDP-43 inclusions (Josephs et al., 2014a; Josephs et al., 2014b) by pharmacologic intervention may lead to a novel therapeutic strategy to slow neuronal death and ameliorate neurologic function in these patients. Further clarification of the sources of cytoplasmic dsRNA may also lead to RNA-based therapeutic strategies. Finally, the development of imaging and CSF biomarkers that detect and predict IFN-I activation are exciting prospective tools to identify patients who may benefit from targeting this pathway within the brain.

## Author Contributions: ( <u>CRediT taxonomy</u>)

Conceptualization, S.R and M.W.A; Methodology, S.R. and M.W.A.; Software, S.R., and M.T., Validation, S.R., B.S., A.S., H.A-L., I.C., A.D.A, L.C., A.C.G.; Formal Analysis,; Investigation, S.R., B.S., A.S., H.A-L., I.C., A.D.A, L.C., A.C.G., M.W.A.; Resources, E.R., M.C., M.P.F; Data Curation, S.R., M.T.; Writing - Original Draft, S.R and M.W.A.; Writing - Review and Editing, all authors; Supervision, P.K.S, B.T.H., and M.W.A; Project Administration, M.W.A.; Funding Acquisition, S.R., M.T., P.K.S., B.T.H. and M.W.A

## Acknowledgements

The authors acknowledge Hannah Brown, Monica Mendelsohn, Jennifer Kirkland for generating the Nd1 mouse line, Sarah Edwards for generating the construct for the Nd2 mouse line. Kristina Holton of the Harvard Medical School Research Computing for assistance with human RNA-seq analysis and use of the Orchestra computing cluster, Clotilde Lagier-Tourenne and Ricardos Tabet for advice about primers for C9, and Richard Axel for critical discussions. The work was supported by the NIH (DP2 OD006662, R21 NS094861 to M.W.A.), P50 AG005134-34 (to B.T.H.), P50-GM107618 (to P.K.S.), Edward R. and Anne G. Lefler Postdoctoral Fellowship, MGH ECOR Postdoctoral Award, and Diversity supplement 5P50AG005134-34 (to S.R.).

## Methods

### Generation and characterization of Nd1 and Nd2 mice

The hAPPsw695 cDNA was amplified using primers with a PacI site flanking the 5’ primer and an AscI and PacI site flanking the 3’ end. For Nd1, the DNA fragment encoding ires-GFP was introduced into the AscI site and the hAPPsw-ires-GFP fragment introduced via a Pac site into pBSRV (Gogos et al., 2000) which contained the tetO sequence followed by an artificial intron and splice site, and an SV40 polyadenylation signal. For Nd2, a DNA fragment encoding ires-mCherry was introduced into the Asc1 site and the hAPPsw-ires-mCherry fragment introduced via a Pac site into pBSRV. These transgenes were injected into the pronuclei of fertilized eggs (Nd1 at Columbia; Nd2 at MGH). Tail DNA from the resulting mice was isolated using standard procedures and analyzed for the transgene by Southern blotting and PCR. Founder mice with the transgene were crossed with OMP-ires-tTA mice (Yu et al., 2004) to determine which founders had active transgenes. For Nd1, compound heterozygote mice of three founders each expressed hAPPsw and GFP in approximately 1% of OSNs and not elsewhere in the brain. For Nd2, compound heterozygote mice of two founders each expressed hAPPsw and mCherry in approximately 1% of OSNs (Nd2) and in approximately 15% of OSNs (CORMAC) (Cao et al., 2012) and not elsewhere in the brain. Founders were backcrossed into C57/BL6 for six generations to generate Nd1 and Nd2.

### RNA isolation and immunoprecipitation of dsRNA

Cytoplasmic RNA was extracted from mouse olfactory epithelium/mouse olfactory bulb/50mg specified region of postmortem human brain using RNeasy kit (Qiagen) except that the standard lysis buffer was exchanged for the lysis buffer RNL [50 mM Tris–HCl, pH 8.0; 140 nM NaCl; 1.5 mM MgCl2; 0.5% (v/v) Nonidet P-40 (1.06 g/ml); and 1 mM DTT added just before use]. Purified RNA was immunoprecipitated using a dsRNA-specific antibody (J2, Scicons) in 200 μl binding buffer (0.025% Triton X-100 in PBS) with 5 μg of J2 antibody and 1 μl RNaseOUT (Life Technologies) rotating overnight at 4°C. J2-bound dsRNA was incubated in binding buffer with 50 μL of prewashed protein-A or G Dynabeads beads for overnight at 4°C, (Life Technologies) followed by 5× washes in cold binding buffer. RNA was then extracted with TRIzol Reagent as above. cDNA was synthesized with SuperScript™ III Reverse Transcriptase (Thermo Fisher). PCR was performed with Phusion High-Fidelity DNA Polymerase (Thermo Fisher) with following primers

#### Primer Name: Primer Sequence

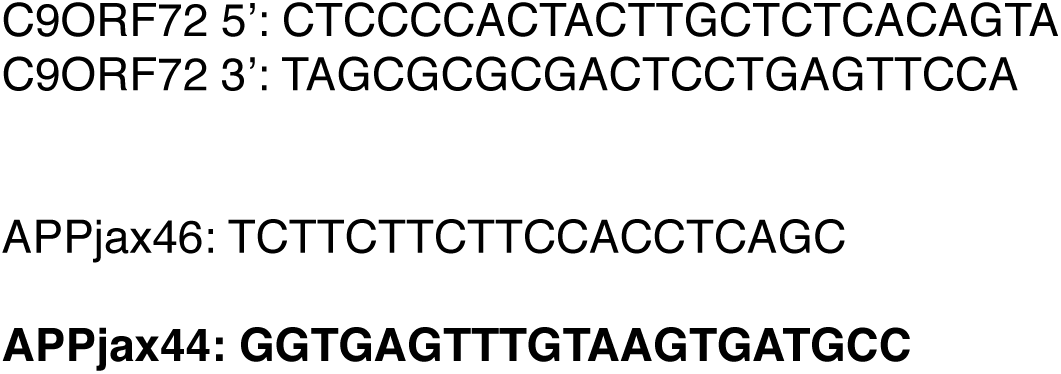

### RNA *in situ* hybridization

RNA *in situ* was performed as previously described (Cao et al., 2012). The templates for cRNA probes were generated by PCR amplification of cDNA libraries synthesized from olfactory epithelia of C57BL/6J mice (stock # 000664, Jackson Laboratories, Bar Harbor, ME), as described below. PCR products were cloned into TOPO pCRII vector (catalog # K4610, Life Technologies, Grand Island NY). Each clone was verified by DNA sequencing (DNA Sequencing Core, Massachusetts General Hospital, Boston, MA). Digoxigenin labeled cRNA probes were synthesized as previously described (Rodriguez et al., 2008) and a 1*μ*l aliquot was run on an 1% agarose gel for quality control.

### cDNA libraries for qPCR

Each olfactory epithelium was ground with an electric homogenizer in 1ml of Trizol reagent (catalog # 15596-018, Life Technologies, Grand Island, NY). RNA was considered of good quality if there was an approximately 2:1 ratio of 28s relative to 18s ribosomal RNA bands on a 1% agarose gel, and a 260/280 spectrophotometer ratio of approximately 2. RNA was DNase treated using the DNA-*free* Kit (catalog # AM1906, Life Technologies, Grand Island NY). 1 *μ*g of purified DNase-free total RNA was used to generate oligodT primed cDNA libraries using the Super Script III First-strand Synthesis System (catalog # 18080-051, Life Technologies, Grand Island, NY). cDNA libraries were cleaned up by ethanol precipitation as follows: samples were diluted to 100*μ*l with water and we added 200*μ*l of a solution consisting of 10% 3M sodium acetate, 95% ethanol, and 1*μ*l of 5 mg/ml Linear Acrylamide (catalog # AM9520, Life Technologies, Grand Island NY). This mixture was placed at -20 degrees for 20 minutes or overnight. The next day the samples were pelleted in a microcentrifuge at 13,500 RPM for 15 minutes, washed 2x with 75% ethanol, air dried for 10 minutes, and resuspended in 100*μ*l of pure water. 1*μ*l of cDNA was used for each 25*μ*l qPCR reaction or for each 25*μ*l PCR amplification to synthesize RNA *in situ* probe templates.

### qPCR

We used the iQ Sybr Green Supermix (catalog # 170-8880, Bio-Rad, Hercules, CA) to perform qPCR in technical triplicates on an iCycler (Bio-Rad, Hercules, CA) with a tm =60 degrees. At least 2-4 biological replicates were performed. qPCR primers were designed using PrimerBank (Spandidos et al., 2010), and were only used for analysis if they had an efficiency of at least 90%.

### Immunofluorescence

For mice 20*μ*m thick cryosections were cut from olfactory epithelia that were fixed and placed onto super frost slides (catalog # 12-550-17, Fisher Scientific, Pittsburg, PA). Sections were dried for 45 minutes at 37 degrees. For sections from animals perfused with 4% PFA we treated the slides for 10 minutes in preheated 10mM citrate buffer pH 6.0 in a food steamer (Oster, Fort Lauderdale, FL). Slides were then incubated for 1 hour with 300-500*μ*l of blocking buffer: 5% non-immune donkey serum, 0.1% triton-X100, in 1xPBS. After this blocking step the slides were dabbed on Kimwipes (catalog # 06-666C, Fisher Scientific, Pittsburg, PA) and we applied 125*μ*l of primary antibody diluted in blocking buffer and incubated in a humidified slide chamber overnight at 4 degrees. The next day the sections were washed 3 times in 1xPBS, then incubated for 2 hours with a 1:500 dilution of Cy3 conjugated donkey anti-mouse secondary antibody (catalog # 706-165-148, Jackson Immunoresearch, West Grove, PA). In cases where amplification was necessary, we used the Cy3 Tyramide Signal Amplification kit (catalog # NEL744001KT, PerkinElmer, Waltham, Massachussetts). For staining human brainsections with the antibody to dsRNA (J2) we cut 8*μ*m thick sections on a microtome, and dried them at 60 degrees for 2 hour. Sections were then deparaffinized in a xylene 2x for 5 minutes and then a graded series of ethanol dips 2 times for 2 minutes each (100%, 95%, 75%, 50%) and then washed in water. Rehydrated slides were then steamed in 0.1M EDTA for 20 minutes and cooled to room temperature. We used the Mouse Elite ABC kit (catalog #PK-6103, Vector Labs, Burlingame, CA) and DAB substrate kit (Catalog # SK-4100, Vector Labs, Burlingame, CA).

### OSN isolation

Freshly dissected olfactory epithelia were minced to ~1mm^3^ cubes with a razor blade on a glass slide, and placed in a scintillation vial for dissociation. To isolate individual cells we used the Papain Dissociation System (catalog # LK003150, Worthington Biochemical Corporation, Lakewood, NJ), with the following modifications: 1) we incubated olfactory epithelia in papain solution for only 15 minutes versus the recommended time without an appreciable loss of cells, 2) after trituration, the cell suspension was filtered through a 40 *μ*m mesh strainer (catalog # 352340, BD Bioscience, San Jose, CA). Dissociated cells were resuspended in 1ml 1x Hank’s balanced salt solution (HBSS) (catalog # 14025, Life Technologies, Grand Island NY) and directly sorted. Each olfactory epithelium was prepared and sorted in 1.5 hours. Samples were visually inspected under a microscope for purity.

### Fluorescence activated cell sorting (FACS)

GFP positive mature OSNs were purified from olfactory epithelia on a FACS Aria II SORPS cell sorter (BD Bioscience, San Jose, CA) with a 70 *μ*m nozzle at a sheath pressure of 70 psi at 5000 events/second. The cells were identified by forward and side scatter and then gated for GFP positive signal relative to a non-GFP expressing control mouse line. The area of GFP positive cells was sorted into a 5ml tube held in a 4 degree chilled chamber containing 500 *μ*l of 1x HBSS. The sorted cells were pelleted at 2,500 rpm for 5 minutes at 4 degrees. The supernatant was discarded and the cells were resuspended in 800 *μ*l of Trizol reagent (catalog # 15596-018, Life Technologies, Grand Island, NY), then frozen on dry ice and maintained at -80 degrees until processed for total RNA isolation. For a quantitative comparison of GFP positive cells in control versus transgenic animals, we compared the percent of GFP positive cells to the entire population of cells in the olfactory epithelium.

### RNA-seq

<u>R</u>NA quality from FACS sorted cells was assessed by an RNA Pico chip on a 2100 Bioanalyzer (Agilent Technologies, Santa Clara, CA) by the Advanced Tissue Resource Center (ATRC, Harvard Neurodiscovery Center, Charlestown, MA). 150 ng of total RNA with a RIN value greater than 8.3 was used to generate sequencing libraries using the Illumina TruSeq mRNA Sample Preparation Kit (catalog # 157894, Illumina, San Diego, CA), as per manufacturers protocol. Four samples, ligated with adapters AR002, AR004, AR006 and AR0012, were pooled and 50bp single end reads were generated from an Illumina HiSeq 2000 by the Next Generation Sequencing Core (Massachusetts General Hospital, Boston, MA). Each sample contained no less than 40 million reads that were mapped to mouse genome mm9. For Nd1 mapped reads were processed for expression analysis using the Cufflinks software pipeline via the Galaxy interface (Blankenberg et al., 2010; Goecks et al., 2010). The gene list was analyzed for molecular pathway signatures using Ingenuity Pathway Analysis software (Qiagen, Redwood City, CA). For RNA-seq analysis of human transcriptomic data from ALS patients with ***C9ORF72*** mutations and controls (GSE67196) we downloaded SRA files containing paired-end reads generated from the cerebellum of patients with ALS bearing the ***C9ORF72*** mutation (SRR1927021, SRR1927023, SRR1927025, SRR1927027, SRR1927029, SRR1927031, SRR1927033) or the cerebellum of healthy patients (SRR1927055, SRR1927057, SRR1927059, SRR1927061, SRR1927063, SRR1927065, SRR1927067, SRR1927069). We performed FastQC and the reads were trimmed with Trimmomatic. StarAligner was used to map the reads to GRCh38 and create a counts table. We then used EdgeR to calculate the differentially expressed genes. Only those genes with a 2-fold change or greater were used in subsequent analysis. Pathway analysis for the human data was done using Enrichr (Chen et al., 2013; Kuleshov et al., 2016).

### Cell counts

A blinded comparison of the number of cleaved-caspase3 or M71 positive cells was done by counting every positive cell on every 10th section though the olfactory epithelium of control and transgenic mice performed in parallel. For gamma-H2AX counts we counted number of nuclei that were pan-positive per olfactory bulb section and normalized it to the area of the olfactory bulb.

### AAV cranial injections

We instilled 2*μ*l of 10^10^ viral genome copies of either AAV9-GFP (catalog # AV-9-ALL854, UPENN Viral Vector Core, PA) or AAV9-FLEX-GFP (catalog #AV-9-PV1963, expressing anti-sense GFP, over the olfactory epithelium ~2mm anterior of the olfactory bulb using a hamilton syringe. Animals were sacrificed 4 weeks after viral instillation for analysis. For quantitation of olfactory epithelium height 5x images were captured from alternating sections of the olfactory epithelium stained for OMP and anti-sense GFP via RNA *in situ* hybridization. On the OMP images we highlighted the boundaries of AAV infection defined by regions positive for anti-sense GFP. We then used a measurement tool in Photoshop to measure the apical-basal distance in pixels of the OMP layer, at approximately equal pixel distance throughout the anti-sense GFP positive layers. We avoided extreme lateral regions within the olfactory epithelium as they naturally undergo more frequent turnover and were likely to confound our data (Vedin et al., 2009).

### ReNcell VM culture

T75 tissue culture flasks were incubated with 5ml of a 1:100 dilution of BD matrigel (catalog #354230, Corning, NY) in ReNcell NSC Maintenance Media (catalog #SCM005, EMD Millipore, Darmstadt, Germany) for one hour at 37°. Prior to adding cells the BD matrigel was aspirated. ReNcell VM neuronal stem cells (catalog #SCC008 Corning, Corning, NY) were maintained in ReNcell NSC Maintenance Media with freshly added 20*μ*g/ml FGF (catalog #03-0002, Stemgent, Lexington, MA) and 20 *μ*g/ml EGF (catalog #GF001, EMD Millipore, Darmstadt, Germany). Once confluent the cells were washed with sterile 1xPBS, and dissociated with Accutase (catalog #A1110501, Invitrogen, Waltham, MA). For differentiation 12,500 cells were seeded onto 96-well plates and maintained in media lacking EGF and FGF. After two weeks the neural stem cells were differentiated into neurons. Differentiated neurons were transfected with polyinosinic-polycytidylic acid (Poly(I:C)) HMW, a dsRNA mimetic (dsRNAmi) (catalog # tlrl-pic, Invivogen, San Diego, California) using Lipofectamine 2000 (catalog #11668027, Invitrogen, Waltham, MA) or Lipofectamine 2000 alone as a control, according to manufacturers protocol. Poly(I:C) conjugated to Rhodamine (catalog # tlrl-picr, Invivogen, San Diego, California) was used to visualize cytoplasmic localization. CellTiter-Glo Luminescent Cell Viability Assay (catalog #G7571, Promega, Madison, WI) was performed according to manufacturers protocol.

### Time-lapse imaging of dsRNAmi

We used ReNcell VM cells that were stably transducer with GFP (Choi et al., 2014; D’Avanzo et al., 2015) and differentiated for 2 weeks were transfected with 2*μ*g/ml of Poly(I:C) conjugated to Rhodamine and imaged on an GE Incell 6000 confocal microscope with controlled humidity and temperature. Cells were imaged every 3 hours for ~2days.

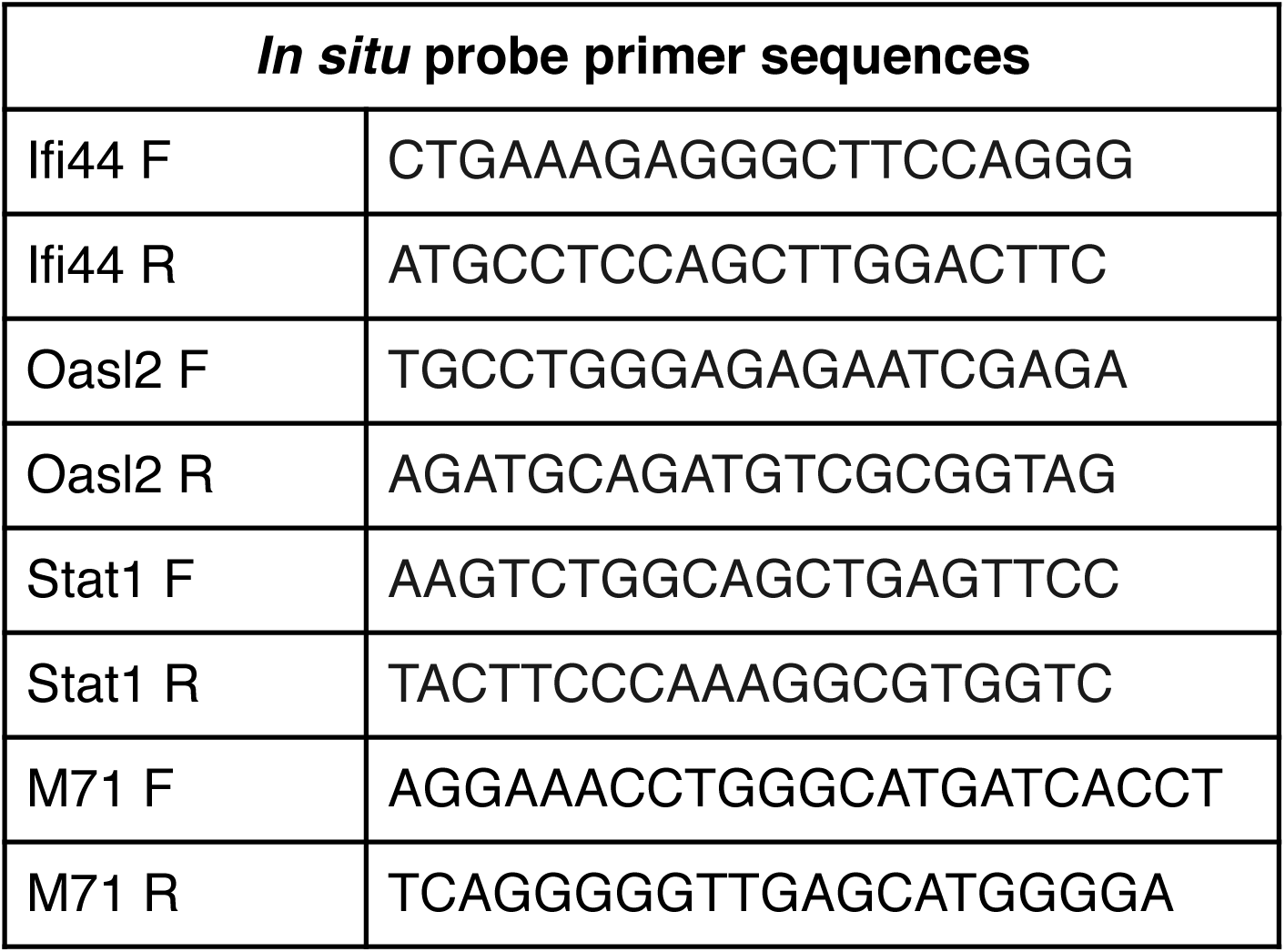

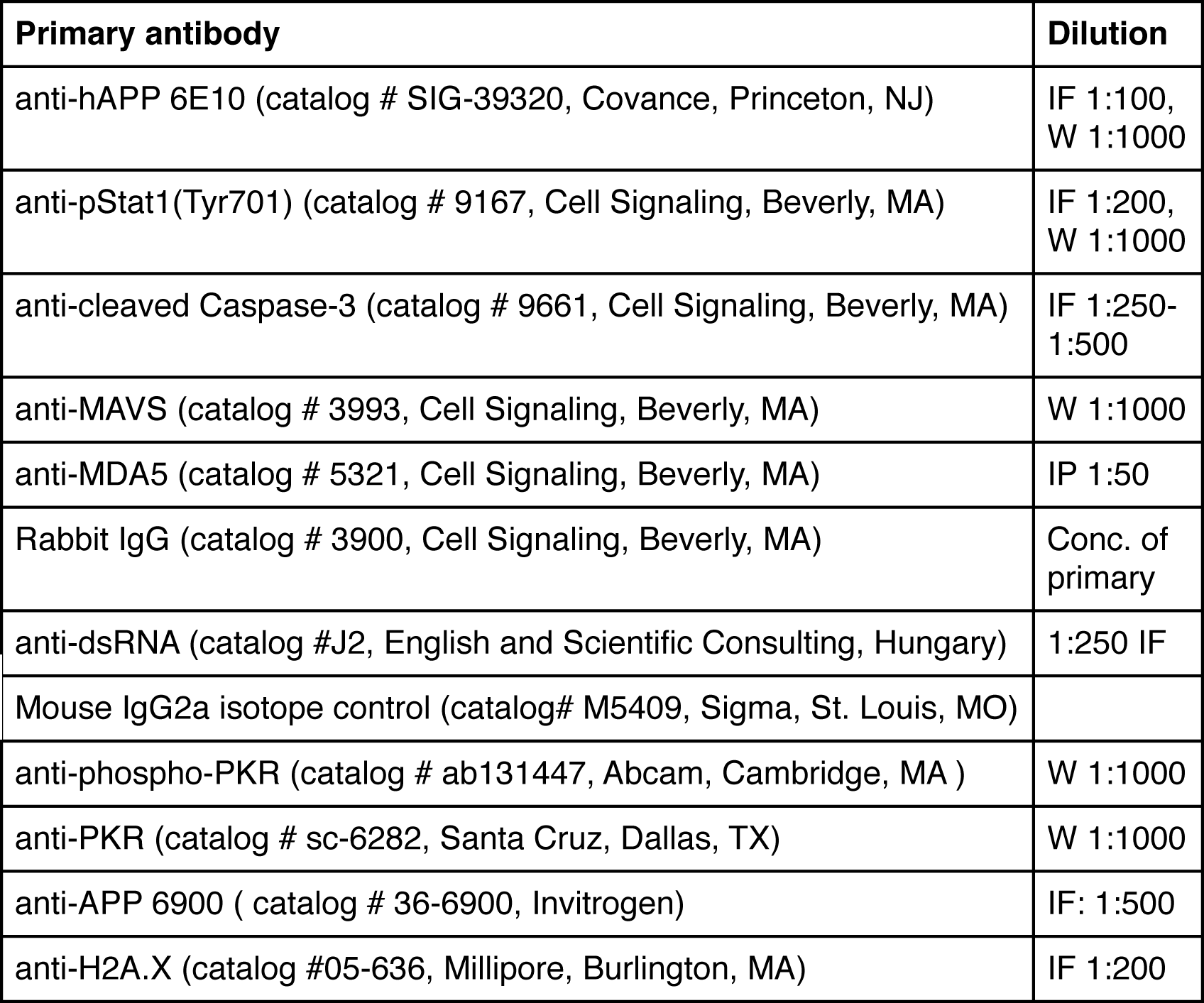

**Supplementary figure 1.**
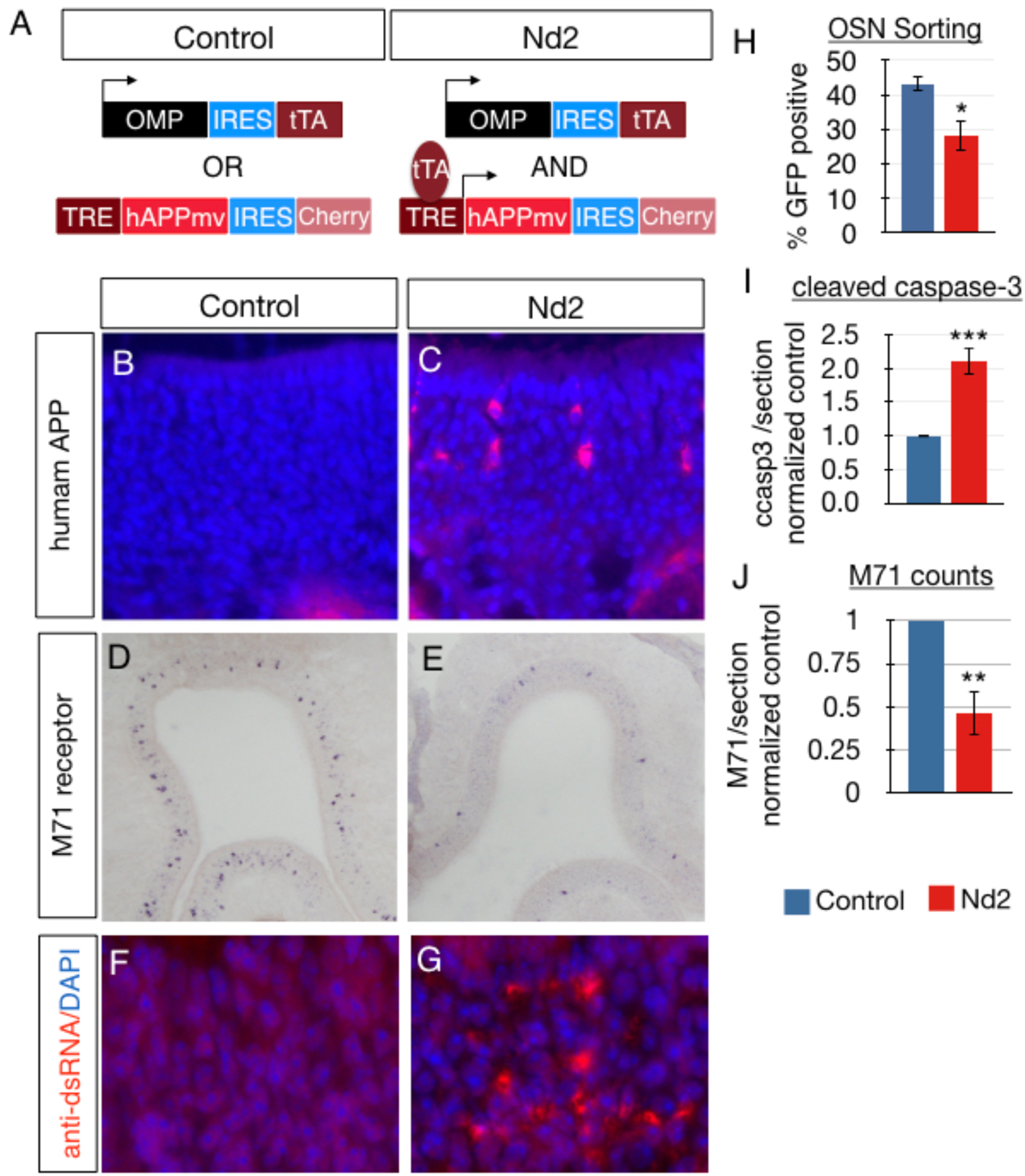
Expression of Nd2 is sufficient to induce mouse olfactory sensory neuron apoptosis. A) A schematic showing the Nd2 transgenic construct; hAPPmv and mCherry are co-expressed with an IRES sequence, which was placed under the transcriptional control of the tetracycline-responsive promoter element (TRE). The tetracycline-controlled transactivator protein is under the control of the endogenous OMP promoter, thereby restricting transgene expression to mature OSNs. We used animals with the transgene alone, or animals expressing tTA alone littermates as controls. B, C) Coronal sections of mouse olfactory epithelia stained by tyramide amplified immunofluorescence with an antibody specific to hAPP (6E10, red) and the nuclear dye, DAPI (blue), in either littermate control (B) or Nd2 transgenic mice (C). D, E) Coronal sections of mouse olfactory epithelia stained by RNA *in situ* hybridization with a digoxigenin labeled probe for the M71 odorant receptor RNA in littermate control (D), or Nd1 mice (E). F, G) Coronal sections of mouse olfactory epithelia stained by tyramide amplified immunofluorescence with an antibody specific to dsRNA (J2) in control (F) or Nd2 mice (G). H) Percentage of olfactory sensory neurons (all mature neurons labeled with GFP) relative to cell in the olfactory epithelium quantified by FACS analysis, in Nd2, p =0.02, n=3. I) Quantitation of the number of cleaved-caspase3 positive cells per section of olfactory epithelium of Nd2, normalized to littermate controls, p= 0.003, n=5. J) Quantitation of the number of M71 positive cells per section of olfactory epithelium in Nd2 normalized to littermate controls, p= 0.0016, n=6

**Supplementary figure 2.**
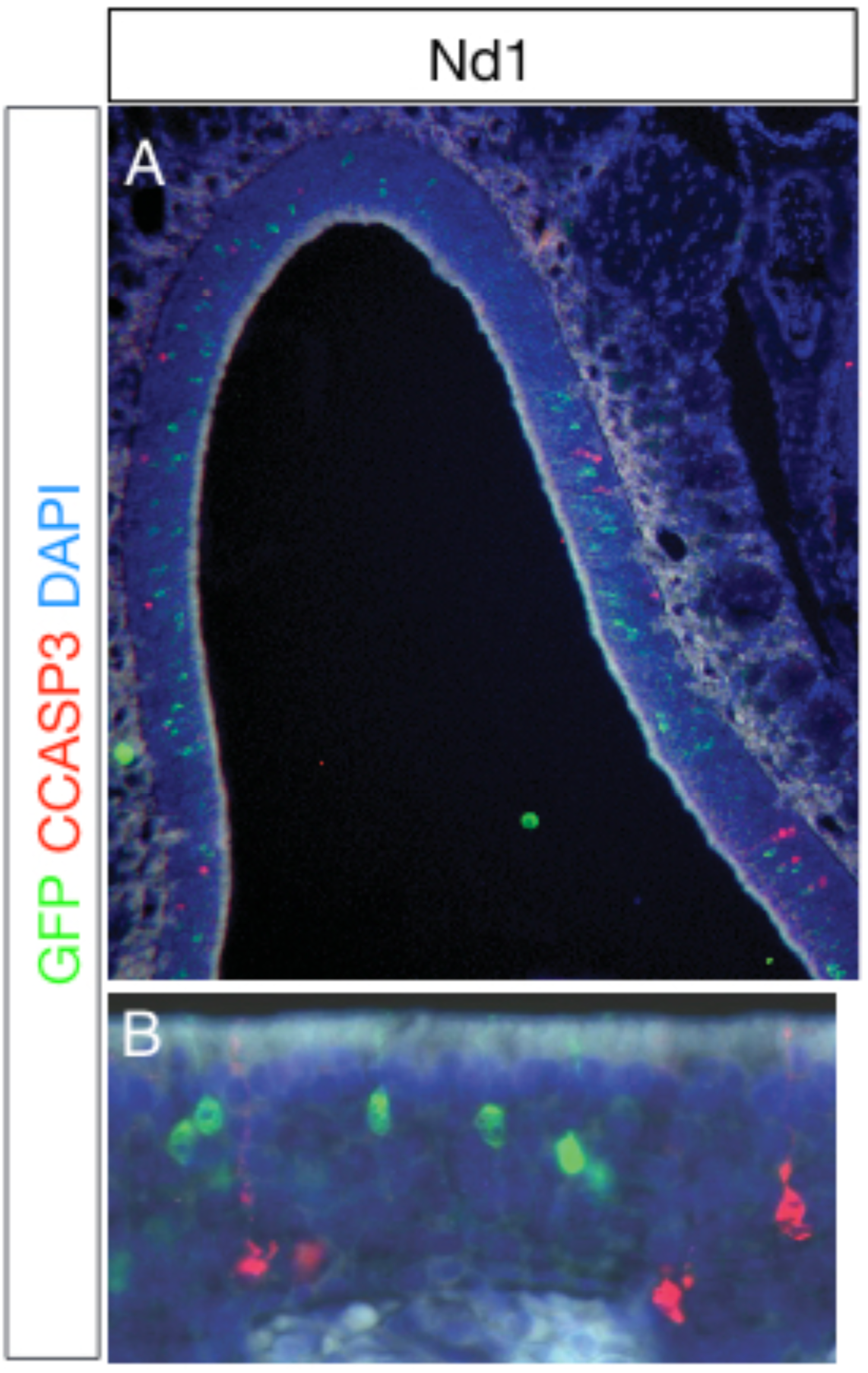
Non-cell-autonomous apoptosis in Nd1 A) Coronal sections of mouse olfactory epithelia co-stained by immunofluorescence with an antibody to cleaved-caspase 3 (red) and GFP (green) in Nd1. B) Quantitation of the number of transgene expressing neurons that undergo apoptosis (p < 0.00001, approximately 1000 CCASP3 positive neurons for each of n=4 mice).

**Supplementary figure 3.**
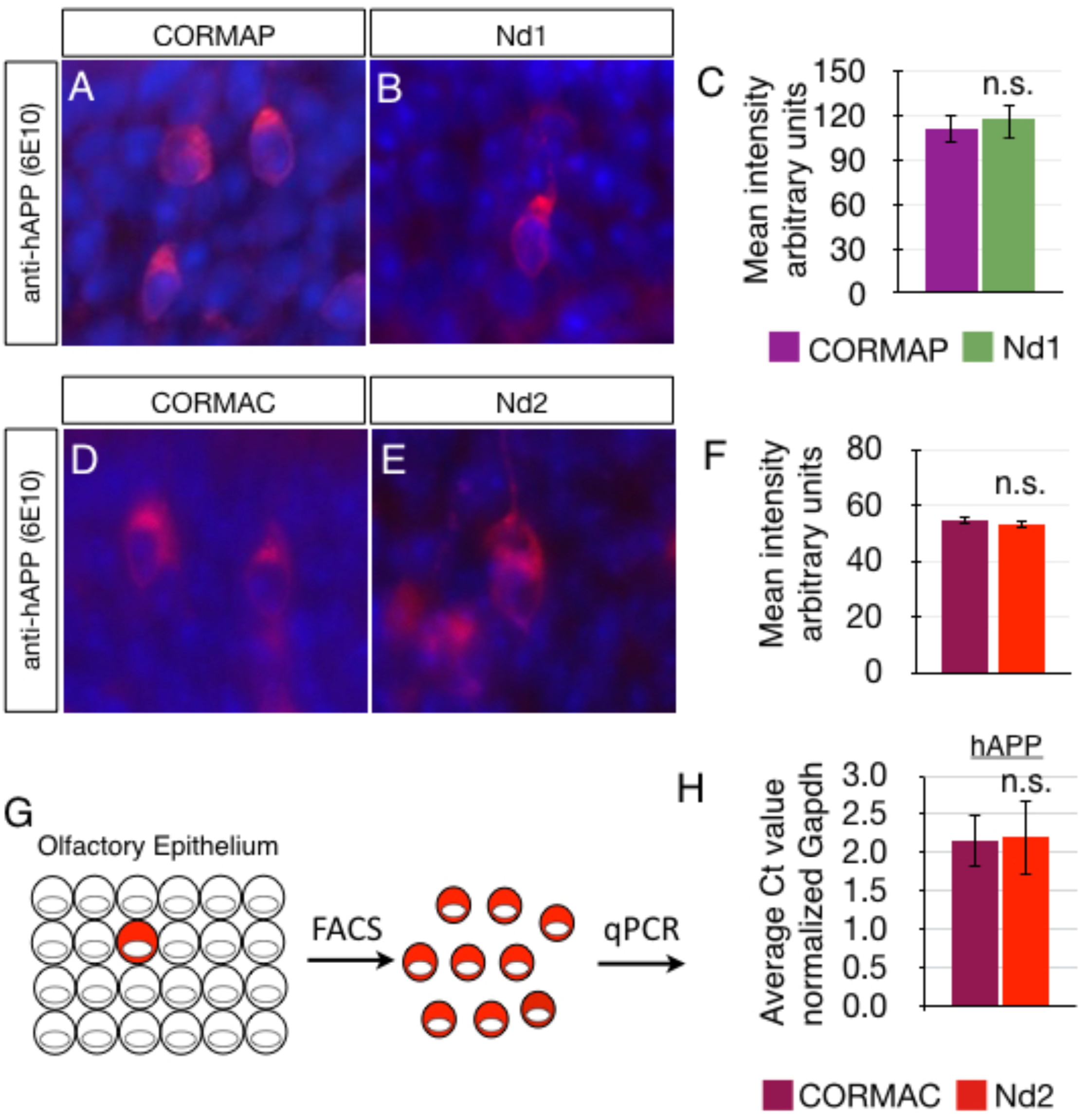
Human-APP expression is equivalent per neuron in CORMAP and Nd1, and CORMAC and Nd2. Coronal sections of fixed mouse olfactory epithelia stained by immunofluorescence with an antibody specific to hAPP (6E10, red) and the nuclear dye, DAPI (blue), in either CORMAP (A), Nd1 (B), CORMAC (D), or Nd2 (E). C) Quantitation of the fluorescence intensity of hAPP expression in CORMAP and Nd1, p=0.620, 100-300 cells in each of n=3 animals. F) Quantitation of the intensity of hAPP expression in CORMAC and Nd2, p=0.72, 100-300 cells in each of n=3 animals. G) Schematic demonstrating that transgene positive cells were isolated from CORMAC and Nd2, which both express mCherry, for quantitation via qPCR. H) Quantitative-PCR analysis of hAPP mRNA levels in FACS isolated transgene positive cells from CORMAC and Nd2 transgenics, p=0.96, n=3 replicates.

**Supplementary figure 4.**
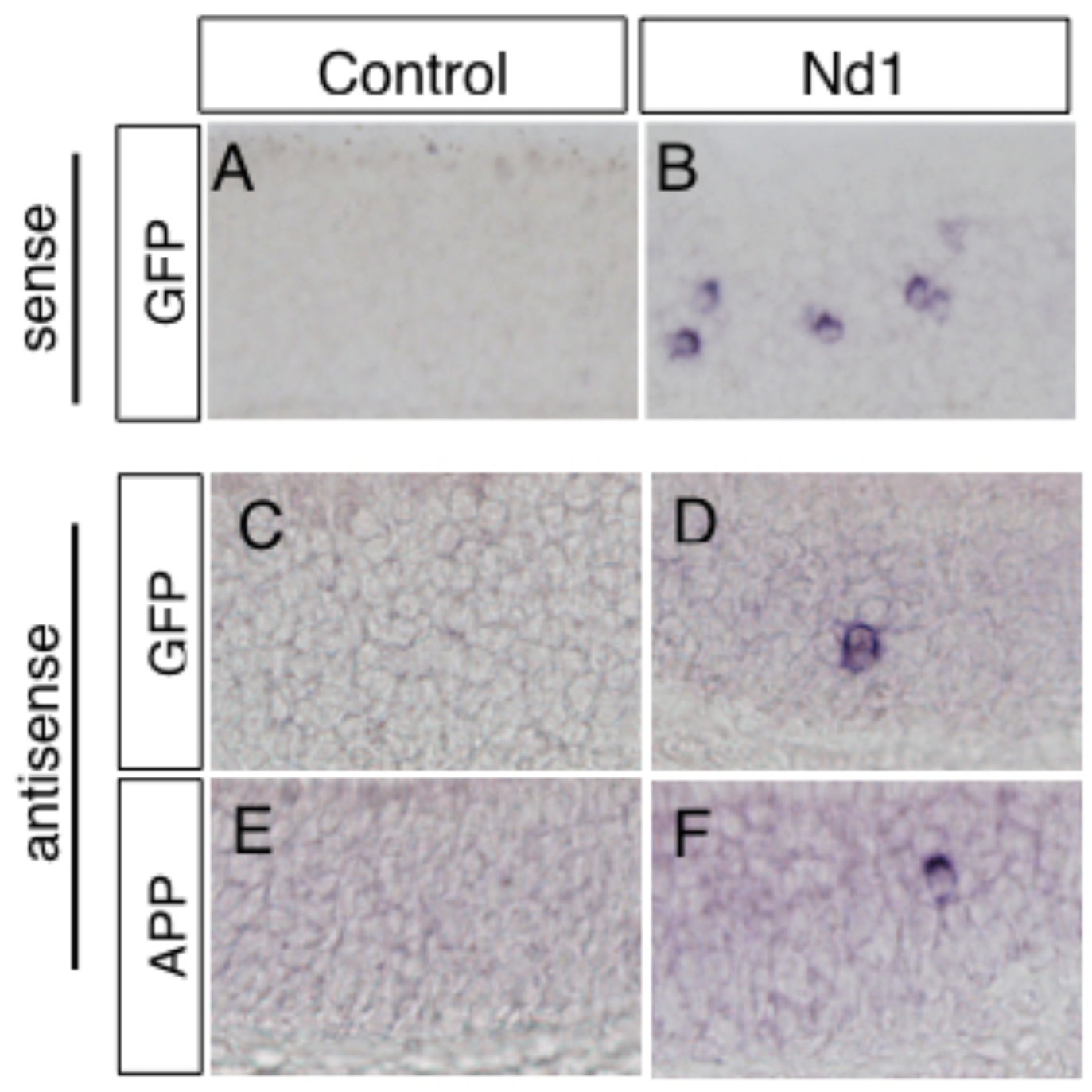
Nd1 transgene produces sense and antisense RNAs. A-F) Coronal sections of mouse olfactory epithelia stained by RNA *in situ* hybridization with digoxigenin lableled probes for: A, B) Sense GFP. C, D) Antisense GFP expression. E, F) antisense APP expression. Note that the APP probe detects both endogenous and transgene derived APP (data not shown) suggesting that only hAPP is generating antisense RNA.

**Supplementary figure 5.**
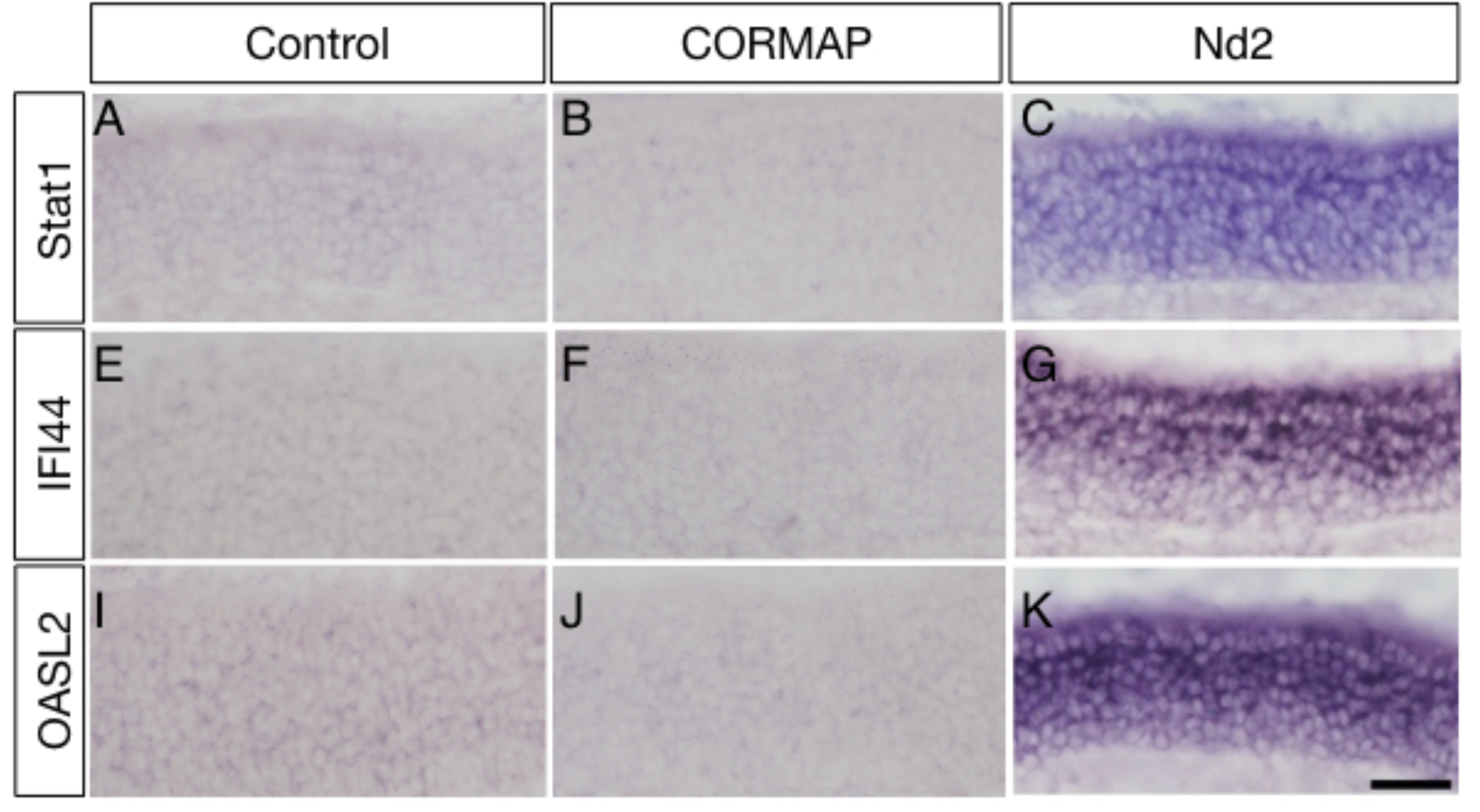
Type I interferon signaling is not unregulated in CORMAP, but is highly unregulated in Nd2 by RNA *in situ* hybridization. Coronal sections of mouse olfactory epithelia stained by RNA *in situ* hybridization with digoxigenin labeled probes for: A-C) Stat1 expression in (A) control, (B) CORMAP, and (Nd2). E-G) IFI44 expression in (E) control, (F) CORMAP, and (G) Nd2. I-K) OASL2 expression in (I) control, (J) CORMAP, and (K) Nd2.

**Supplementary figure 6.**
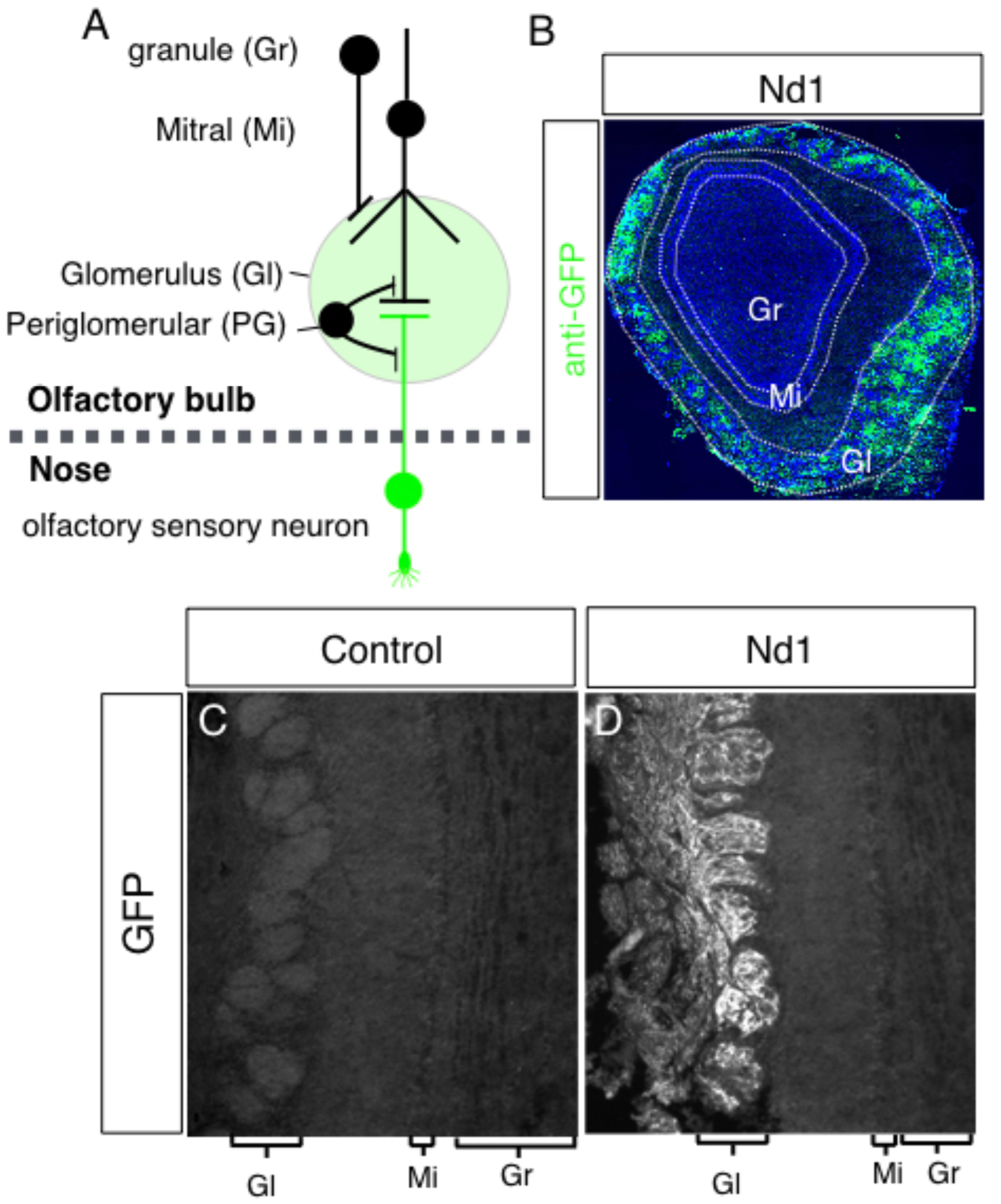
Nd1 transgene expression in the olfactory bulb is restricted to olfactory sensory neurons axons. A) Schematic showing the major neuronal cell types in the olfactory bulb. The green color represents GFP positive axons from OSNs in the neuropil of the olfactory bulb (Gl). B) Immunofluorescence staining of the olfactory bulb in Nd1 for GFP. Note that expression of GFP in the olfactory bulb is restricted to neuropil in the olfactory bulb, and is not expressed the mitral cell (Mi) or granule cell (Gr) layers. C,D) higher magnification (20x) view showing GFP is restricted to axons in Gl layer, near the surface of the olfactory bulb (i.e. not internal to the olfactory bulb).

**Supplementary figure 7.**
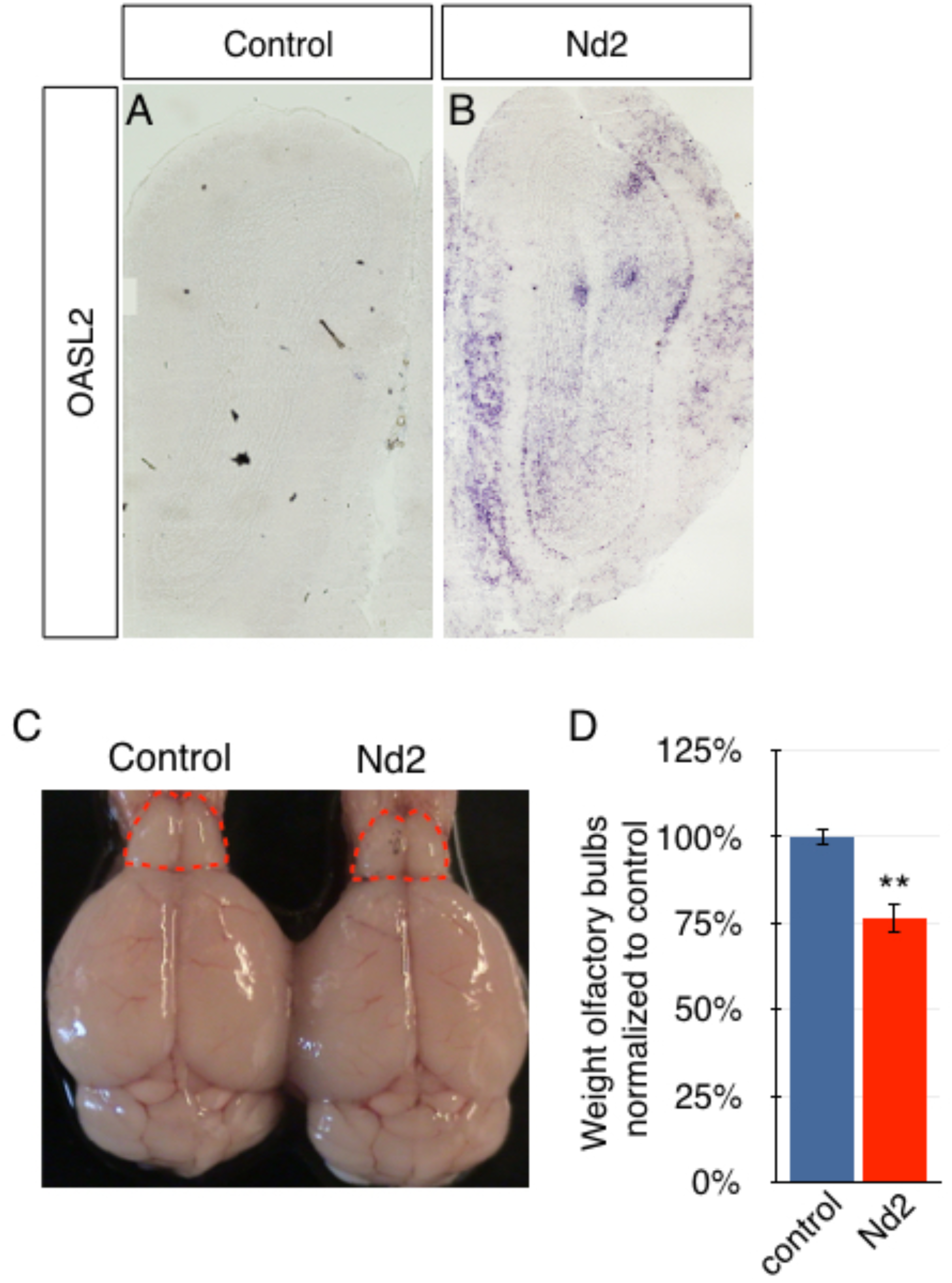
Propagation of IFN-I signaling and neurodegeneration in Nd2. A,B) Coronal sections of mouse olfactory epithelia stained by RNA *in situ* hybridization with a digoxigenin labeled probe for OASL2, as a marker for IFN-I signaling (we see similar results with RNA *in situ* probes for IFI44 and Stat1, data not shown). C) Olfactory bulbs in Nd2 mice are smaller than littermate control mice. D) Quantitation of olfactory bulb weights in Nd2 and littermate control mice, n=3, p=0.008.

**Supplementary figure 8.**
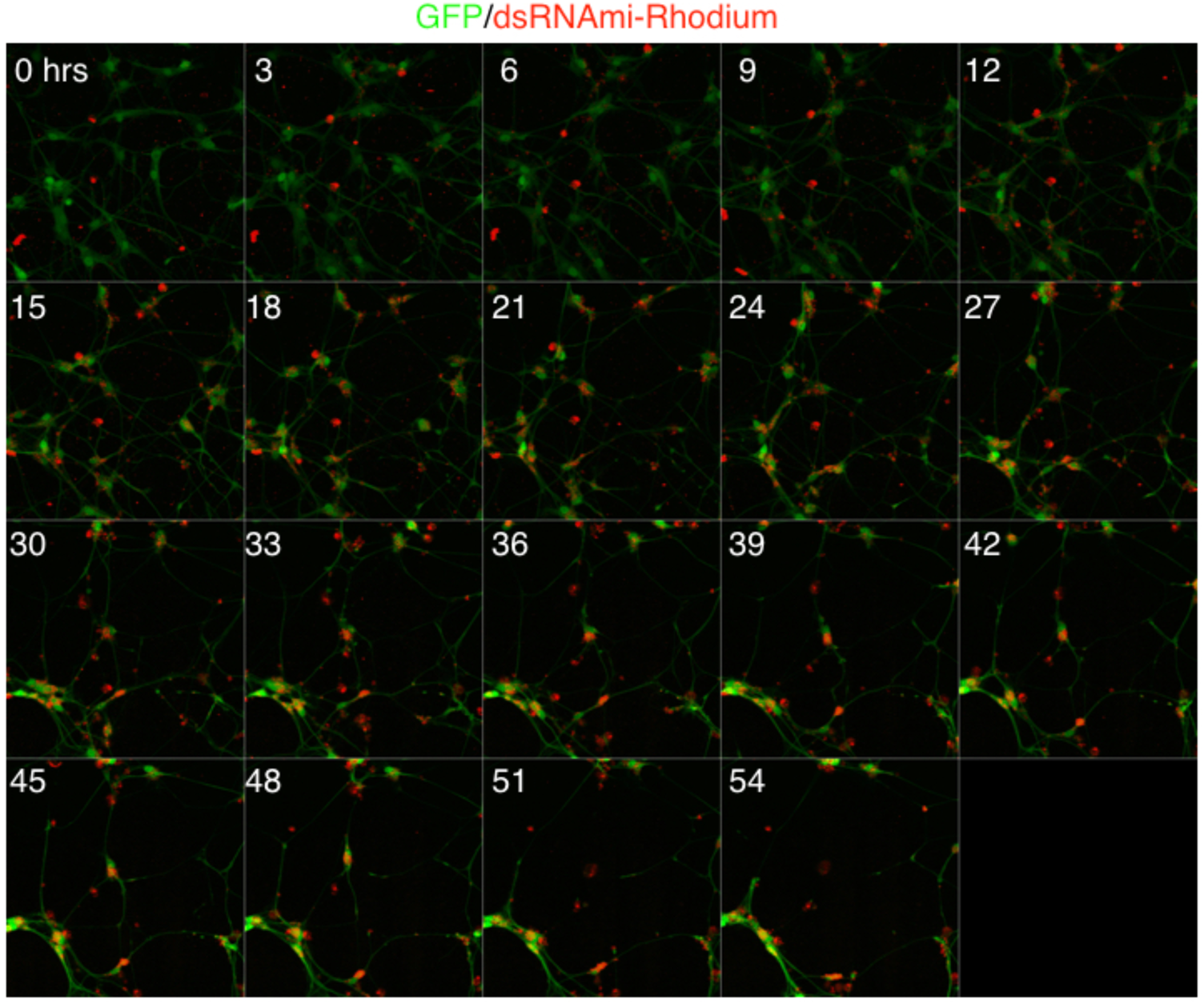
DsRNAmi localizes to the perinuclear region of the cytoplasm leading to neuronal death. Montage of Supplementary Movie 1. Every image contains the hour post transfection of GFP-positive neurons with Rhodamine-labeled dsRNAmi. (20x).

**Supplementary figure 9.**
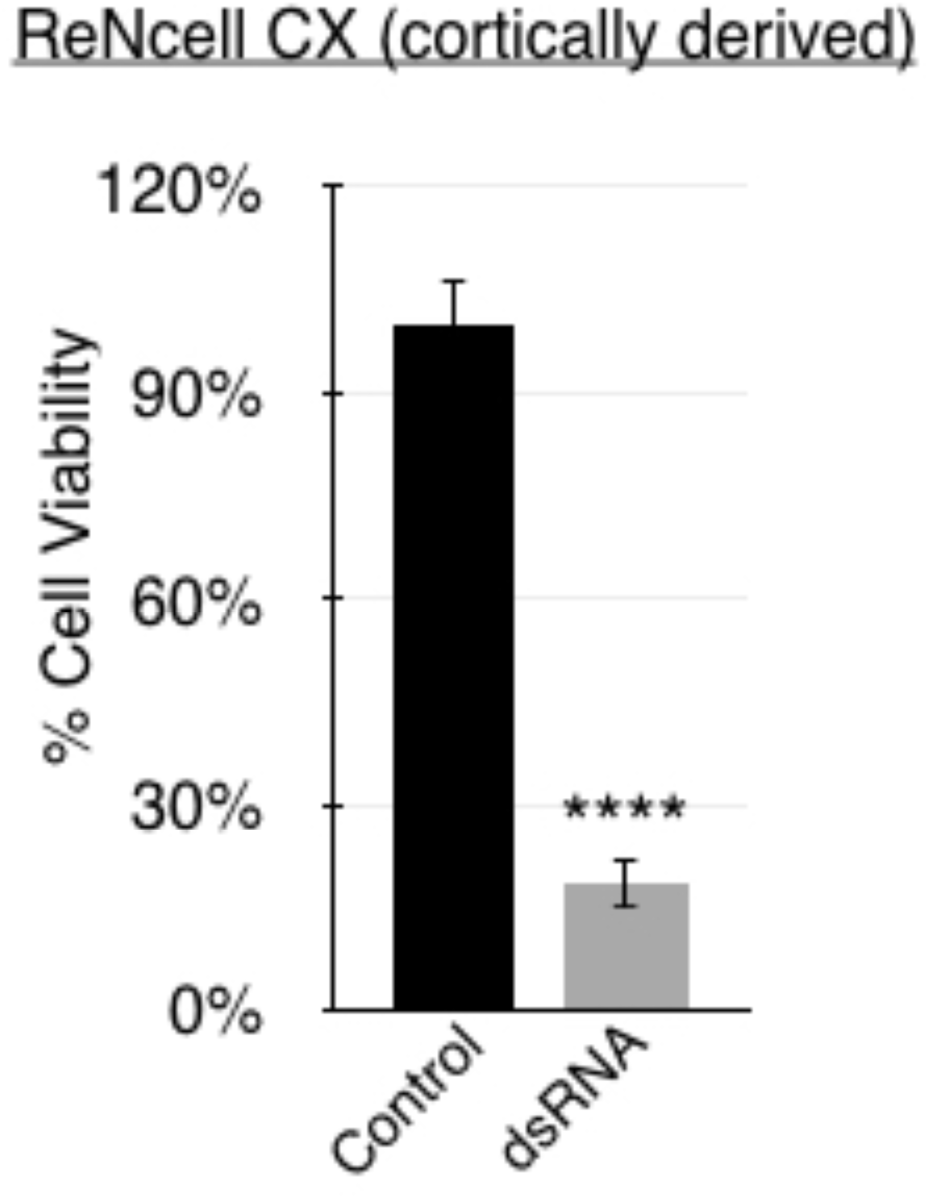
DsRNA is sufficient to induce neuronal cell death in differentiated human cortical neurons. ReNcell CX (cortical neurons) were differentiated for two weeks and then transfected with 2 *μ*g/mL of dsRNAmi. The percent viability of cells was determined after two days using CellTiter-Glo reagent to measure ATP levels.

**Supplementary figure 10.**
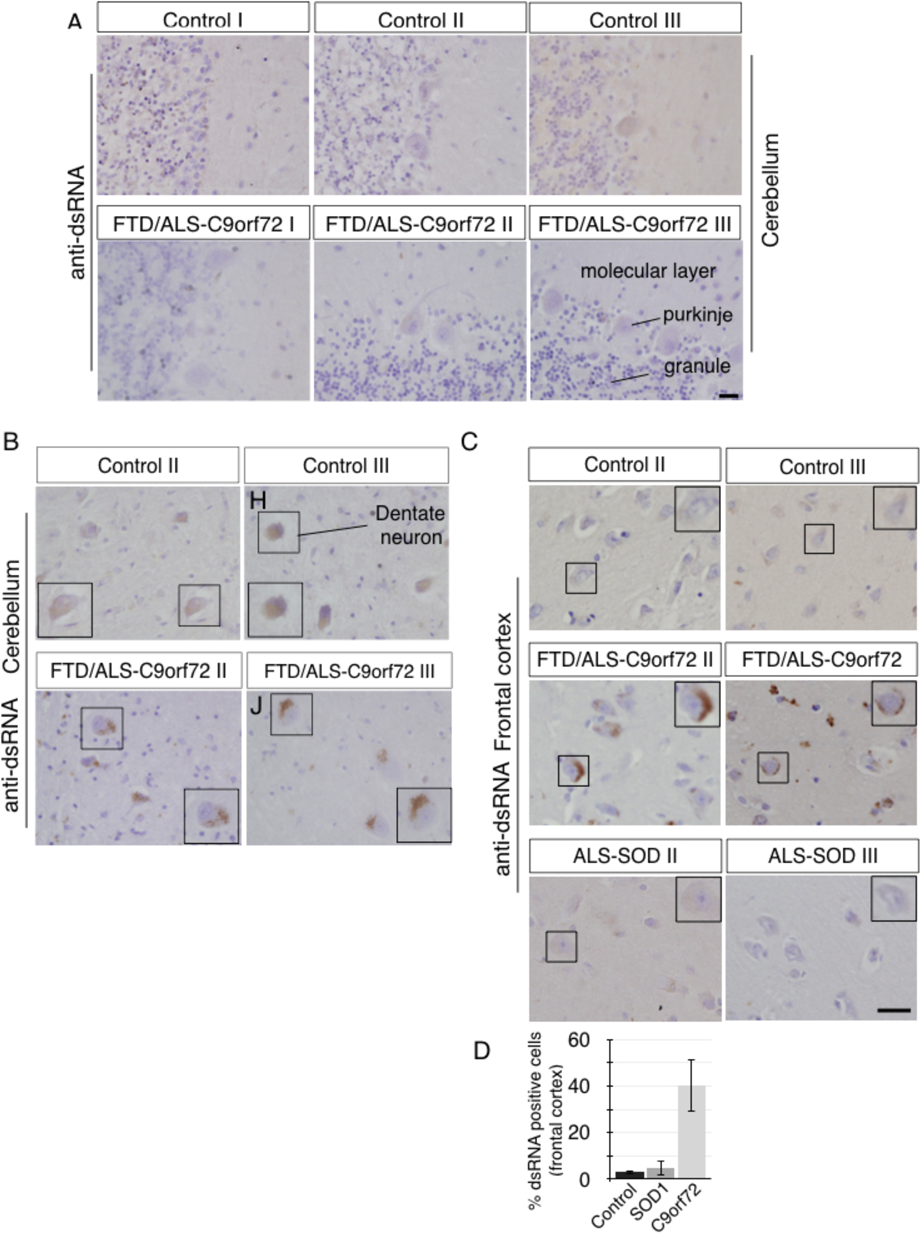
DsRNA expression in additional clinical cases. A) DsRNA immunohistochemistry with antibody specific to dsRNA (J2) on fixed thin sections of cerebellum from normal control brains or on brain sections from ALS/FTD patients that contained the ***C9ORF72*** mutation showing lack of discernable dsRNA in the molecular, Purkinje, or granule layer of the cerebellum of ALS/FTD patients with the ***C9ORF72*** mutation. B) DsRNA is detected at higher levels in the dentate neurons of the cerebellum in ***C9ORF72*** (I-J) relative to controls (G-H). C) Quantitation of the number of dsRNA positive cells in the frontal cortex from control, SOD1 mutation carriers (p= 0.54, n=3) and ***C9ORF72*** mutation carriers (p=0.004, n=3).

**Supplementary figure 11.**
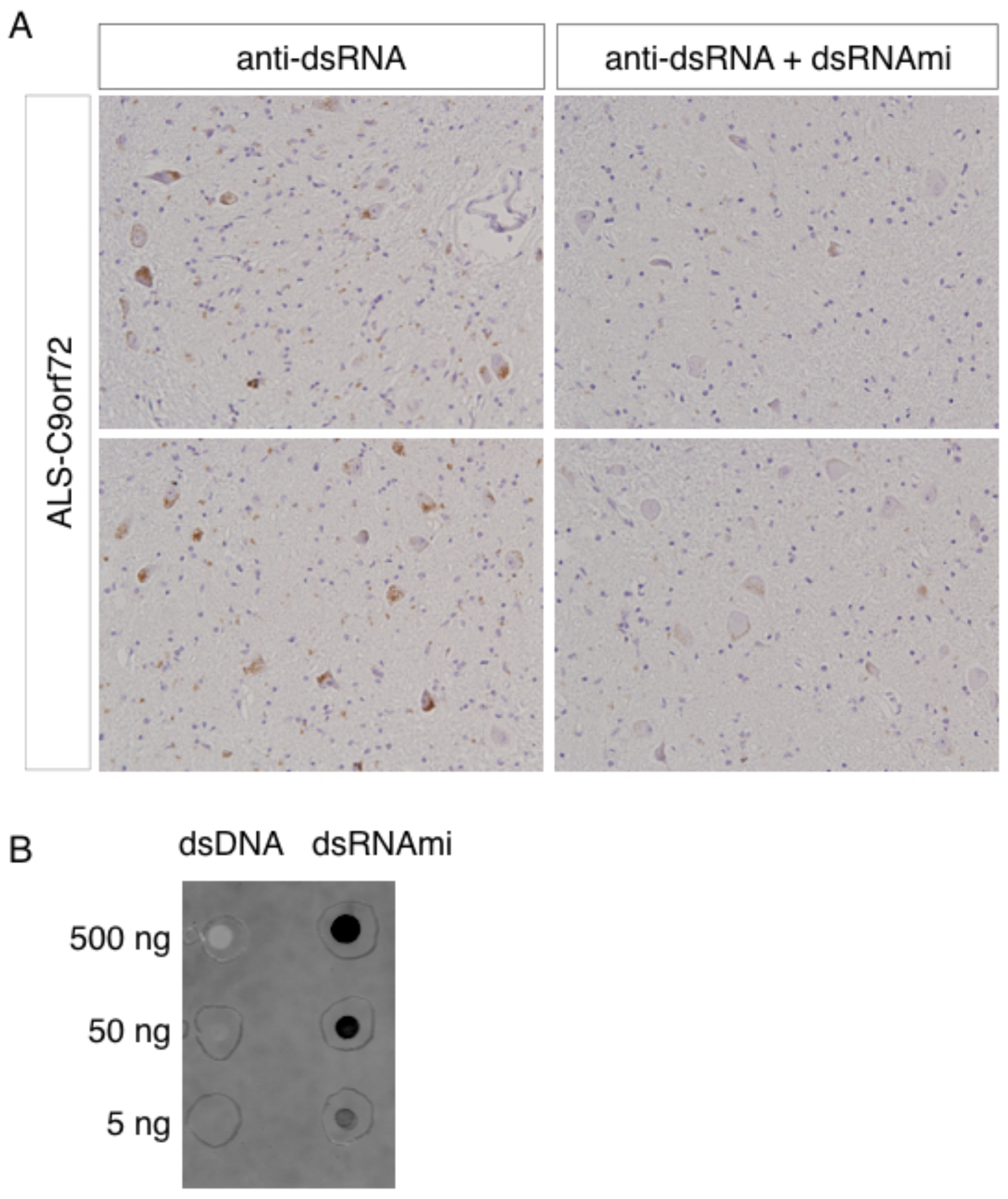
Immunohistochemistry with antibody specific to dsRNA (J2) on fixed thin sections is specific to dsRNA. Anti-dsRNA antibody (J2) was added directly to tissue sections (A,C) or pre-incubated with dsRNAmi prior to application onto tissue sections (B,D). E) A membrane was blotted with either dsDNA or dsRNAmi at the same concentration. The anti-dsRNA antibody (J2) binds specifically to the dsRNAmi, but not the dsDNA.

**Supplementary table 1.** Nd1 transgenes, but not CORMAP transgenes, underwent complex genetic rearrangements. Sanger sequencing confirmation of whole genome sequencing of jumping libraries of Nd1 and CORMAP showing that transgenic sequence of Nd1 but not CORMAP underwent a complex genetic rearrangement.

**Supplementary table 2.** RNA-seq analysis of RNA from FACS isolated OSNs in Nd1 compared to littermate controls.

**Supplementary table 3.** GSEA analysis of differentially expressed genes in Nd1 are enriched in genes that correspond to Interferon/virally stimulated cells and cancer/ genomic instability regulated gene sets.

|  |  | Control | Nd1 |
| --- | --- | --- | --- |
| Interferon $\alpha$ | Ifnab | 0 | 0 |
|  | Ifna1 | 0 | 0 |
|  | Ifna2 | 0 | 0 |
|  | Ifna4 | 0 | 0.416 |
|  | Ifna5 | 0 | 0 |
|  | Ifna6 | 0 | 0 |
|  | Ifna7 | 0 | 0 |
|  | Ifna9 | 0 | 0 |
|  | Ifna11 | 0 | 0 |
|  | Ifna12 | 0 | 0.037 |
|  | Ifna13 | 0 | 0 |
|  | Ifna14 | 0 | 0 |
| $\beta$ | Ifnb1 | 0 | 0 |
| $\gamma$ | Ifng | 0 | 0 |

**Supplementary table 4.** Interferon-α is the only interferon expressed by neurons in Nd1 transgenic mice. RNA-seq FPKM values from isolated mature olfactory sensory neurons in Nd1 transgenic and control mice.

**Supplementary table 5.** Pathway analysis of human C9orf72 patients- GO Molecular Function term enrichment

**Supplementary table 6.** Pathway analysis of human C9orf72 patients- Lincs L1000 Ligand Perturbation enrichment

